# *Arabidopsis* CONSERVED BINDING OF EIF4E1 negatively regulates the NADPH oxidase RESPIRATORY BURST OXIDASE HOMOLOG D

**DOI:** 10.1101/2022.01.27.478037

**Authors:** Jeoffrey George, Martin Stegmann, Jacqueline Monaghan, Cyril Zipfel

**Author notes:** Center for Plant Cell Biology, Department of Botany and Plant Sciences, University of California, Riverside, Riverside, CA, USA. Phytopathology, TUM School of Life Sciences, Technical University of Munich, Freising, Germany. Department of Biology, Queen’s University, Kingston K7L 3N6, Ontario, Canada (Jacqueline Monaghan).

## Abstract

Cell-surface pattern recognition receptors sense invading pathogens by binding microbial or endogenous elicitors to activate plant immunity. These responses are under tight control to avoid excessive or untimely activation of cellular responses, which may otherwise be detrimental to host cells. How this fine-tuning is accomplished is an area of active study. We previously described a suppressor screen that identified *Arabidopsis thaliana* mutants with regained immune signaling in the immunodeficient genetic background *bak1-5*, which we named *modifier of bak1-5* (*mob)* mutants. Here, we report that *bak1-5 mob7* restores elicitor-induced signaling. Using a combination of map-based cloning and whole-genome resequencing, we identified *MOB7* as *CONSERVED BINDING OF EIF4E1* (*CBE1*), a plant-specific protein that interacts with highly-conserved eukaryotic translation initiation factor eIF4E1. Our data demonstrate that CBE1 regulate the accumulation of RESPIRATORY BURST OXIDASE HOMOLOG D (RBOHD), the NADPH oxidase responsible for elicitor-induced apoplast reactive oxygen species (ROS) production. Furthermore, several mRNA decapping and translation initiation factors co-localize with CBE1 and similarly regulate immune signaling. This study thus identifies a novel regulator of immune signaling and provides new insights into ROS regulation, and more generally translational control during plant stress responses.

## Introduction

The restriction of invading organisms is governed by passive and active defenses, which are effective against all types of plant pathogens and pests, including viruses, insects, nematodes, and parasitic plants^1^. On the cell surface, conserved microbial molecules called pathogen- or microbe-associated molecular patterns (PAMPs/MAMPs) or plant-derived damage-associated molecular patterns and phytocytokines (hereafter, generally referred to as elicitors) are recognized by pattern recognition receptors (PRRs)^2,3^. For example, in *Arabidopsis thaliana* (hereafter Arabidopsis), the PRRs FLAGELLIN SENSING 2 (FLS2), EF-TU RECEPTOR (EFR), and PEP1 RECEPTOR 1 (PEPR1) and PEPR2 recognize bacterial flagellin (cognate ligand, flg22), bacterial EF-Tu (cognate ligand, elf18), and the endogenous Atpep1 (and related peptides), respectively^4–6^. These PRRs interact with the common co-receptor BRASSINOSTEROID INSENSITIVE 1-ASSOCIATED KINASE 1 (BAK1) in a ligand-dependent manner^7–9^. Following heterodimerization, numerous cell signaling events are initiated, including activation of receptor-like cytoplasmic kinases (RLCKs), production of apoplastic reactive oxygen species (ROS) by the NADPH oxidase RESPIRATORY BURST OXIDASE HOMOLOG D (RBOHD), altered ion fluxes, activation of calcium-dependent protein kinases (CDPKs), mitogen-activated protein kinase (MAPK) cascades, callose deposition, and large-scale transcriptional programming^10,11^.To maintain immune homeostasis, plants use multiple strategies to adjust the amplitude and duration of immune responses^11^. These include limiting the ability of PRRs to recruit their cognate co-receptors, regulation of signaling initiation and amplitude at the level of PRR complexes (*i.e*. post-translational modifications, protein turn-over), monitoring of cytoplasmic signal-transducing pathways, and control of transcriptional reprogramming^11^.

To identify loci involved in plant immunity, a forward genetic screen was previously conducted in the immunodeficient mutant *bak1-5*, called the *modifier of bak1-5* (*mob*) screen^12^. This EMS-induced suppressor screen of *bak1-5* phenotypes identified 10 mutants in nine allelic groups, named *mob1* to *mob10*, with partially restored elicitor-induced ROS production^12–14^. Through this suppressor screen, novel regulators of immune signaling have been discovered. *MOB1* and *MOB2* encode CALCIUM-DEPENDENT PROTEIN KINASE 28 (CPK28), which negatively regulates immune signaling by controlling the accumulation of the RLCK BOTRYTIS-INDUCED KINASE 1 (BIK1), a central kinase involved in immune signaling downstream of multiple PRRs^12,15,16^. *MOB4* encodes CONSTITUTIVE ACTIVE DEFENSE 1 (CAD1)^13^. CAD1 is involved in immunity at different levels by controlling programmed cell death and regulating the homeostasis of the phyllosphere microbial community^17,18^. MOB6 corresponds to SITE-1 PROTEASE (S1P), which controls the maturation of the endogenous RAPID ALKALINIZATION FACTOR 23 (RALF23) peptide to regulate immune signaling via the receptor kinase FERONIA^14,19,20^. Hence, we predict that the identification of remaining *MOB* genes will continue to unravel mechanisms of immune regulation.

Here, we report that MOB7 corresponds to CONSERVED BINDING OF EIF4E1 (CBE1), a plant-specific protein that associates with the 5’ mRNA cap^21^ and the translation initiation factor eIF4E1^22^. We show that CBE1 co-localizes with ribonucleoprotein complexes, and that *cbe1* and other translational regulator mutants display enhanced accumulation of RBOHD protein, resulting in enhanced anti-bacterial immunity and ROS production, possibly through translational control of RBOHD protein levels.

## Results

### The *mob7* mutation rescues *bak1-5* immuno-deficiency

In the present study, we describe and characterize the *mob7* mutation. First, we confirmed that the *mob7* mutation was maintained in the M_5_ generation as *bak1-5 mob7* mutants displayed restored ROS production in seedlings upon treatment with the elicitors elf18 and flg22 (Figure 1A). In addition, the *mob7* mutation restored ROS production in adult leaves upon elicitation with elf18, Atpep1, and chitin; however, no difference was observed with flg22 (Figure 1B; Figure S1A-D). Despite rescuing the ROS phenotype quantitatively, *mob7* did not restore the delayed peak of ROS burst observed in *bak1-5* (Figure S1A-E). A late immune output triggered by several elicitors is the inhibition of seedling growth^10^. While seedling growth inhibition is largely blocked in *bak1-5* mutants^9,23^, it was restored in *bak1-5 mob7* upon prolonged exposure with elf18, flg22, or Atpep1, while mock-treated seedlings grew similar to wild-type (WT) Col-0 (Figure 1C; Figure S1F). Furthermore, immunity to the hypovirulent bacterial strain *Pseudomonas syringae* pathovar *tomato* (*Pto*) DC3000 *COR*^*-*^ was restored in *bak1-5 mob7* compared to *bak1-5* (Figure 1D). Altogether, these results show that *mob7* partially restores immunity in *bak1-5*.

**Figure 1.**
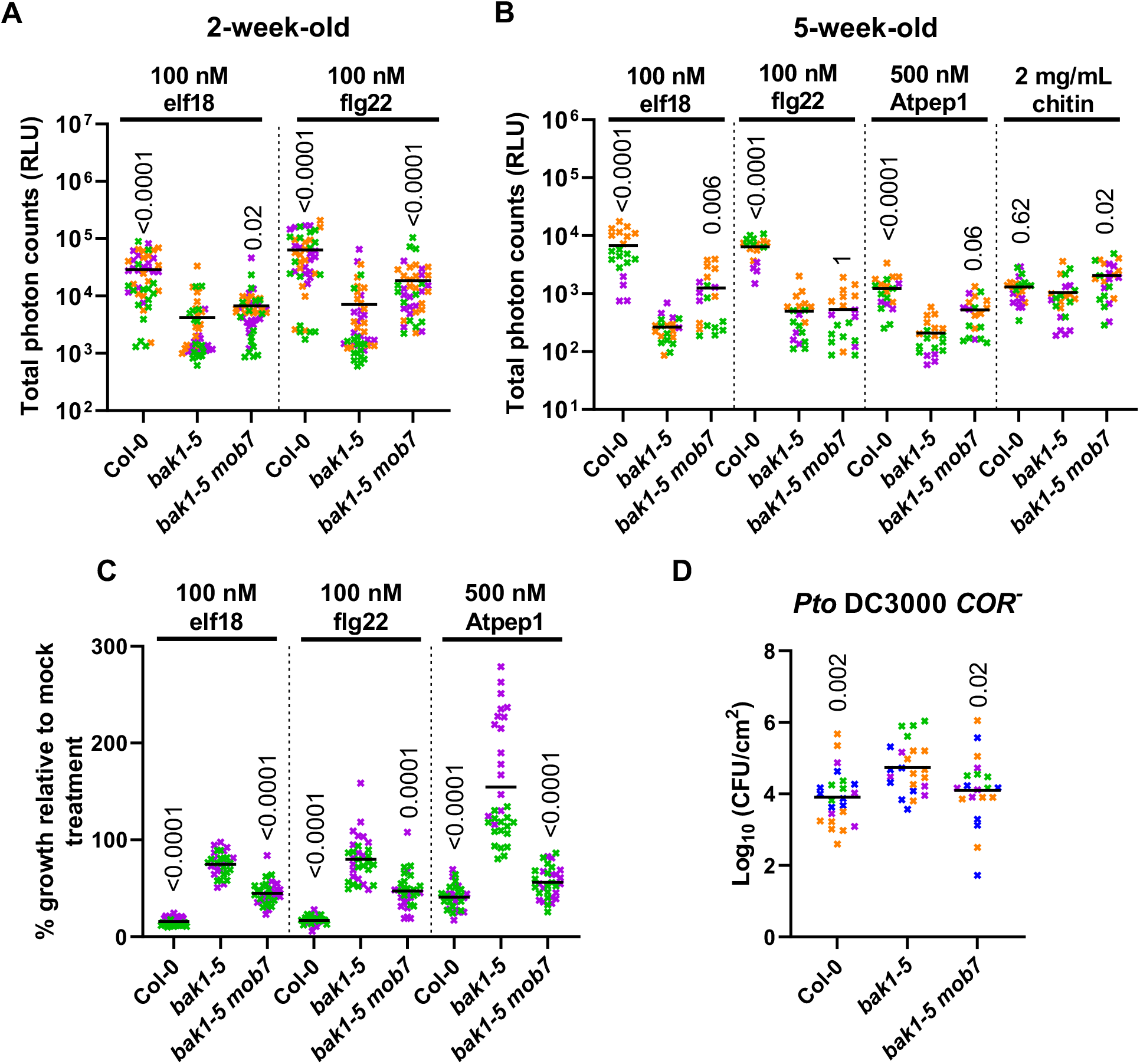
*mob7* restores immune signalling in *bak1-5*. (A-B) Total ROS accumulation measured as relative light units (RLU) over 60 min recording after treatment with the corresponding elicitors on (A) 2-week-old seedlings (n=12-16) or (B) leaf discs from 5-week-old leaves (n=4-8). Horizontal lines represent the means from 3 independent experiments (n=4-8) (C) Growth inhibition is represented as relative fresh weight compared to untreated seedlings in response to the indicated elicitors. Horizontal lines represent the means from 2 independent experiments (n=12-17). (D) Bacterial growth (colony-forming unit -CFU /cm^2^) in leaves spray-inoculated with 10^7^ CFU/mL (OD_600_ = 0.2) *P. syringae* pv. *tomato* (Pto) DC3000 *COR*^*-*^ and sampled at 3 dpi. Horizontal lines represent the means from 4 independent experiments (n=4-8). (A-D) Symbol colors indicate different experiments. Numbers above symbols are p-values from (A, B, C) Dunn’s or (D) Dunnett’s multiple comparison test between corresponding genotypes and *bak1-5*.

### Identification of *MOB7* as *CBE1*

Using the elicitor-induced ROS phenotype of *mob7* and map-based cloning of the F_2_ population from the outcross of *bak1-5 mob7* (Col-0 ecotype) with L*er*-0, linkage analysis revealed 3 regions of interest (Figure S2). Whole-genome resequencing of bulked F_2:3_ segregants that rescued seedling growth inhibition with 1 μM Atpep1 identified a single nucleotide polymorphism in *AT4G01290*, a gene that encodes CONSERVED BINDING OF EIF4E1 (CBE1) (Figure 2A). The G to A transition is located at the last nucleotide of the second exon (Figure 2B; Figure S3A), which leads to a premature stop codon resulting in reduced *CBE1* expression and the production of a truncated protein (Figure 2C; Figure S3A). It is possible that the premature stop codon in *mob7* is recognized by the nonsense-mediated mRNA decay (NMD) machinery, which links premature translation termination to mRNA degradation^24^. Knock-down alleles from independent T-DNA insertions in *CBE1* phenocopied the increased elf18-induced ROS production and normal growth observed in *mob7* single mutant (Figure 3A; Figure S3A-D), while WT segregants from these T-DNA lines have the same phenotype as Col-0 (Figure 3A; Figure S3A-C). We thus feel confident that the mutation we identified in *CBE1* explains the *mob7* phenotypes.

**Figure 2.**
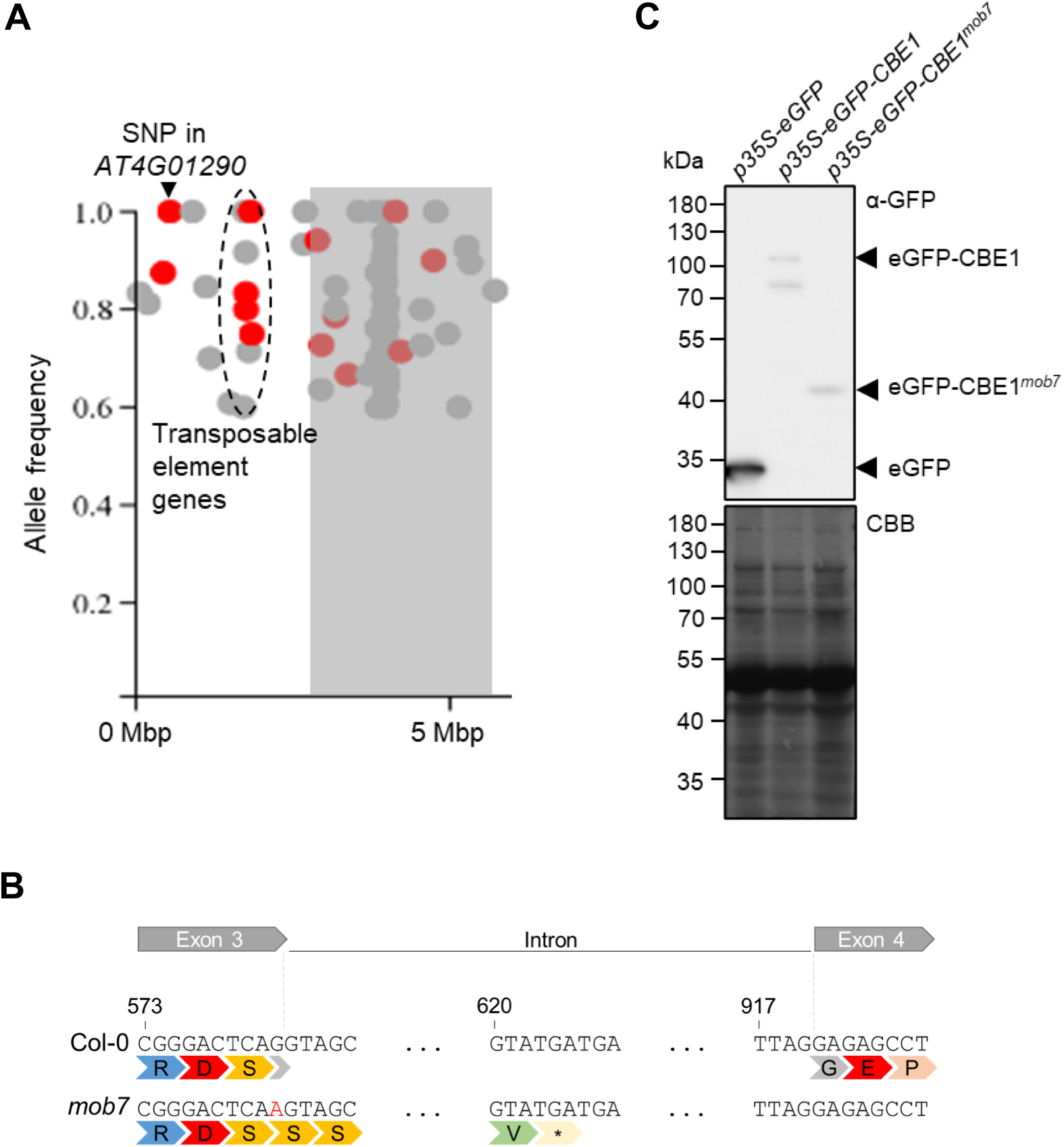
*mob7* mutation maps to *CONSERVED BINDING OF EIF4E1* resulting in a truncated protein. (A) Density plot of SNPs at the top arm of chromosome 4 using CandiSNP software (Etherington *et al*., 2014). SNPs with an allele frequency below 60% were removed from the plots. Non-synonymous SNPs are shown in red and others in grey. Grey rectangles indicate the centromere. The dashed area delimits several non-synonymous SNPs in transposable element genes. Mbp, mega base pairs. (B) The *mob7* mutation leads to a premature stop codon within the intron downstream of exon 3. The top symbols delimit nucleotides from exons 3, 4 and intron within *AT4G01290*. The number indicates the nucleotide position relative to the adenosine of the start codon. The second line shows amino acids corresponding to codons above. The EMS-induced SNP in *mob7* is indicated in red. Star indicates a stop codon. (C) Immunoblot analysis using anti-GFP after transient expression in *N. benthamiana*. Coomassie Brilliant Blue (CBB) stain is shown as loading control. Experiment was repeated once with similar results.

**Figure 3.**
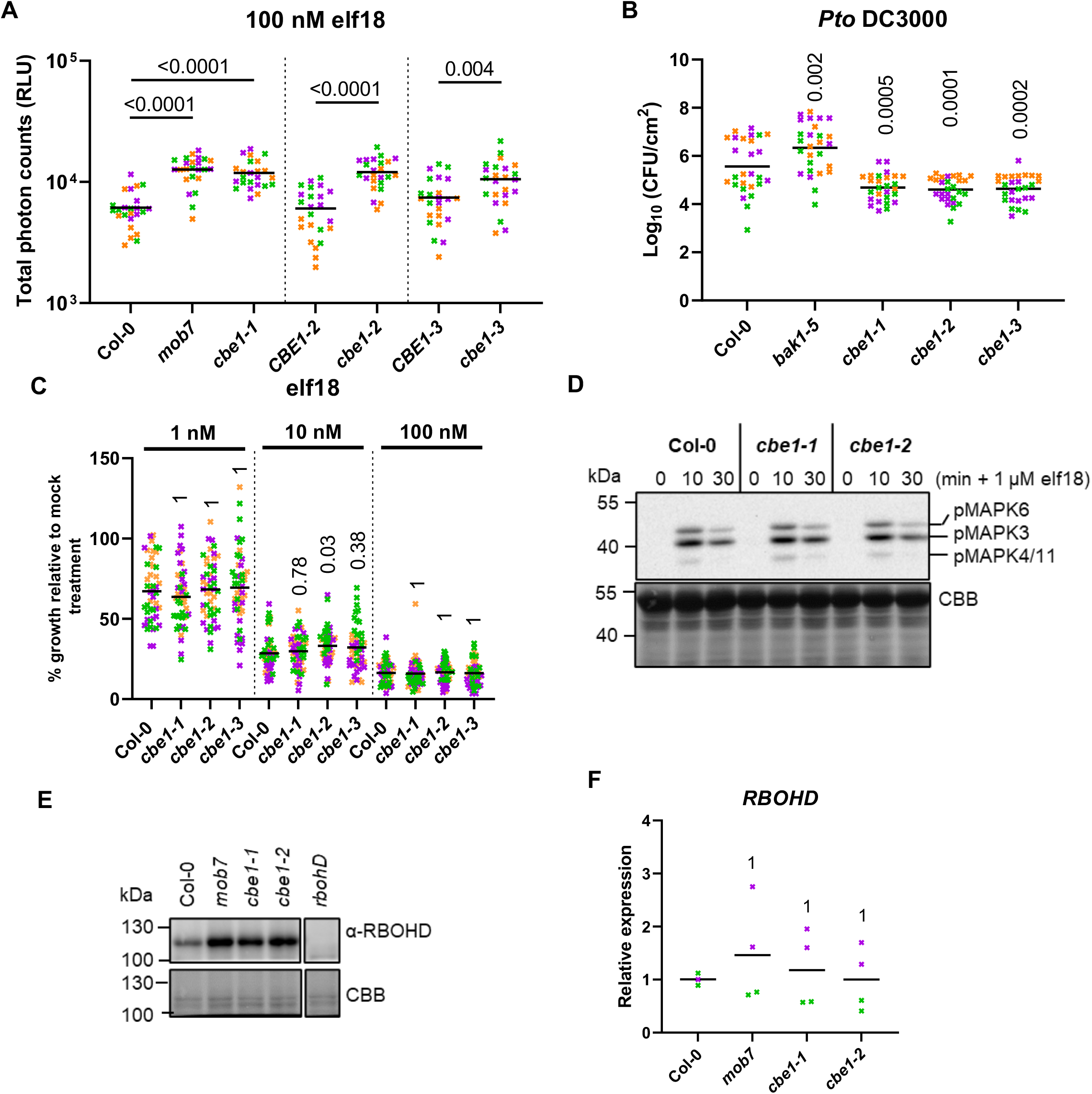
CBE1 negatively regulates elicitor-induced ROS production and RBOHD protein levels. (A) Total ROS accumulation measured as RLU over 60 min recording after treating leaf discs from 5-week-old plants with 100 nM elf18. Horizontal lines represent the means from 3 independent experiments (n=8). (B) Bacterial growth (CFU/cm^2^) in leaves spray inoculated with 10^7^ CFU/mL (OD_600_= 0.2) *P. syringae* pv. *tomato* DC3000 and sampled at 3 dpi. Horizontal lines represent the means from 3 independent experiments (n=9). (C) Growth inhibition represented as percentage of fresh weight in response to 1, 10 or 100 nM elf18 relative to mock treated seedlings. Horizontal lines represent the means from 3 independent experiments (n=16). (D) Immunoblot analysis of elf18-induced MAPK phosphorylation using anti-phospho-p44/42 in leaf discs from 5-week-old Arabidopsis leaves treated with 1 μM elf18 for the indicated time. Coomassie Brilliant Blue (CBB) stain is shown as loading control. Experiment was repeated twice with similar results. (E) Immunoblot analysis of RBOHD (anti-RBOHD) and BAK1 (anti-BAK1) protein accumulations in 5-week-old Arabidopsis leaves from corresponding genotypes. Coomassie Brilliant Blue (CBB) stain is shown as loading control. Experiment was repeated twice with similar results. (F) qRT-PCR of *RBOHD* transcripts in corresponding genotypes. Expression values relative to the *U-BOX* housekeeping gene are shown. Horizontal lines represent the means from 2 independent experiments (n=1-2). Numbers above symbols are p-values from (A, B) Dunnett’s or (C, F) Dunn’s multiple comparison test between corresponding genotype and *bak1-5*.

### CBE1 is a negative regulator of elicitor-induced ROS production and immunity

While mutation of *CBE1* results in increased ROS production induced by various elicitors (Figure 3A; Figure S4A), and enhanced immunity to *Pto* DC3000 (Figure 3B), we did not observe any differences in seedling growth inhibition or MAPK activation between different *cbe1* alleles and Col-0 (Figure 3C,D; Figure S4B). Given the apparent specific impact of *cbe1* mutations on ROS production, we tested whether transcripts and/or protein levels for the NADPH oxidase RBOHD were affected. Interestingly, while no significant reproducible difference could be observed at the transcript level (Figure 3F: ref. 22), RBOHD protein accumulation was consistently higher in *cbe1* mutants (Figure 3E). These results indicate that CBE1 regulates RBOHD post-transcriptionally or co/post-translationally, which could thus explain the effect on ROS production and immunity.

### CBE1 co-localizes with ribonucleoprotein complexes

As CBE1 is known to interact with the translation initiation factors eIF4E and eIFiso4E^22^, which localize to ribonucleoprotein complexes associated with the 5’ cap of mRNA transcripts, we were interested to investigate the subcellular localization of CBE1. When transiently expressed in *Nicotiana benthamiana*, CBE1-GFP displays a nucleo-cytoplasmic subcellular distribution, additionally localizing to distinct cytoplasmic foci (Figure 4A). Comparatively, while CBE1^*mob7*^-GFP similarly localizes to the cytoplasm and nucleus, localization in cytoplasmic foci is not apparent (Figure 4B). To investigate the localization of CBE1 within cytoplasmic foci, colocalization was measured using Pearson correlation coefficient with different ribonucleoprotein complex markers^25^. Active translation is located within polysomes while processing bodies (P-bodies) and stress granules are generally associated with decay and storage of mRNA, respectively^26^. To differentiate those different sub-complexes, we used marker proteins. Associated with P-bodies, DECAPPING 1 (DCP1)^27^ is a member of the decapping complex, which is responsible for removal of the 5’ cap, while UP-FRAMESHIFT SUPPRESSOR 1 (UPF1)^28^ is a factor of NMD. Although generally associated with active translation within polysomes, the translation initiation factor eIF4E^29^ and POLY(A) BINDING PROTEIN 2 (PAB2)^30^ also localize to stress granules, together with the RNA binding proteins OLIGOURIDYLATE BINDING PROTEIN 1B (UBP1B)^30^ and RNA BINDING PROTEIN 47C (RBP47C)^29^. We observed the highest co-localization correlation between CBE1 and DCP1 as well as partial co-localization between CBE1 and UPF1 (Figure 4C; Figure S5). To a lesser extent, CBE1 also co-localized with polysomes and stress granule markers eIF4E, UBP1B, RBP47C, and PAB2 (Figure 4C; Figure S5). This indicates that CBE1 co-localizes with ribonucleoprotein complexes and suggests a role for CBE1 in P-bodies.

**Figure 4.**
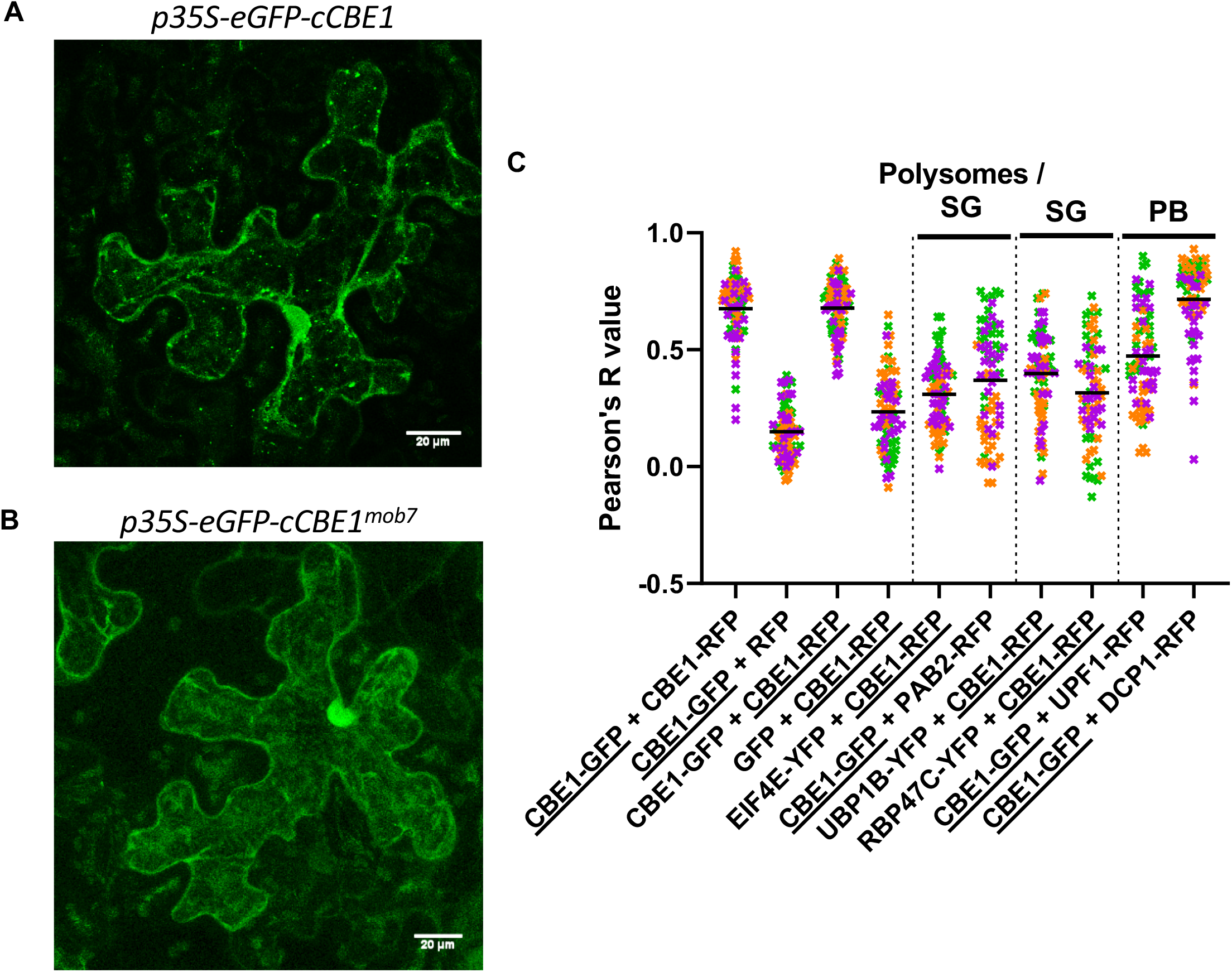
CBE1 localizes predominantly to processing bodies among ribonucleoprotein complexes. (A, B) Confocal images of CBE1-GFP (A) or CBE1^*mob7*^-GFP (B) after transient expression in *N. benthamiana*. Each picture is a z-stack projection. The scale bar corresponds to 20 μm. (C) Quantitative co-localization analysis for CBE1 with polysomes / stress granules (SG), SG specific and P-bodies (PB) markers after transient co-expression in *N. benthamiana*. The Pearson correlation coefficient (R) was calculated with five ROIs (25 μm^2^) per image (n=5, images) and the proteins underlined refer to the channel used to draw the ROIs.

### RBOHD accumulation is affected in mutants of additional translation factors

We next tested if RBOHD accumulation and subsequent immune outputs are affected in mutants lacking components of the translation initiation complex (*i.e*. eIF4E1, eIFiso4E, eIF4G, eIFiso4G1/2)^31^, or P-bodies (*i.e*. PAT1)^32^. As PAT1 was shown to be guarded by the nucleotide-binding site leucine-rich repeat receptor (NLR) SUPPRESSOR OF MKK1 MKK2 2 (SUMM2)^32^, the double mutant *pat1-1 summ2-8* was also analyzed together with the single mutants *pat1-1* and *summ2-8*. Similar to *cbe1-1, eif4e1* and *pat1* mutants, and to a lesser extent *eif4g*, showed a similar ROS phenotype upon elicitor treatment (Figure 5A). Accordingly, *eif4e1* and *pat1-1* mutants also displayed an increase in RBOHD protein level similar to *cbe1* (Figure 5B,C), suggesting that RBOHD may be regulated by these factors.

**Figure 5.**
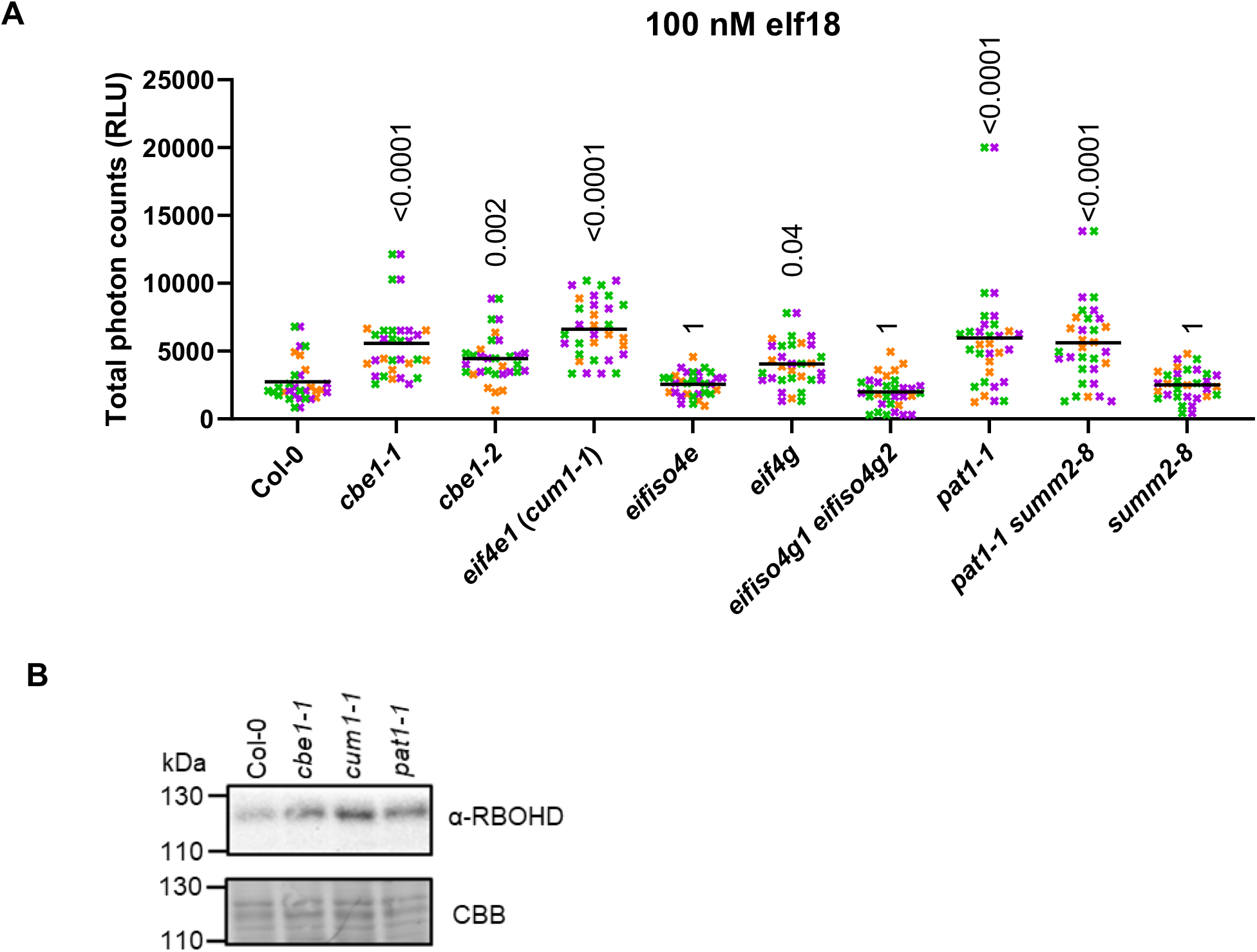
Translation factor eIF4E and decapping factor PAT1 also play a role in ROS production. (A) Total ROS accumulation measured as RLU over 60 min recording after treatment with 100 nM elf18 on leaf discs from 5-week-old plants: Horizontal lines represent the means from 3 independent experiments (n=8-12). The symbol colors indicate the different experiments. Numbers above symbols are p-values from Dunn’s multiple comparison test between the corresponding genotypes and Col-0. (B) Immunoblot analysis of RBOHD (anti-RBOHD) protein accumulations in 5-week-old Arabidopsis leaves from the corresponding genotypes. Coomassie Brilliant Blue (CBB) stain is shown as loading control. Experiment was repeated twice with similar results.

## Discussion

Immune signaling relies on tight regulation to allow a proportionate and timely response^11,33^. Here, we report that CBE1 contributes to RBOHD protein accumulation and consequently elicitor-induced ROS production and anti-microbial immunity. Similarly, mutants of the decapping factor PAT1 and the translation initiation factor eIF4E phenocopied *cbe1*. Overall, this suggests that CBE1, PAT1, and eIF4E regulate RBOHD levels, either post-transcriptionally or co/post-translationally, and thereby affect elicitor-induced ROS production. Translational regulation of plant immunity has recently been proposed, as elicitor perception induces global translational reprogramming^34–36^ and remodeling of the cellular RNA-binding proteome^37^. Notably, some of these RNA-binding proteins (RBPs) control transcripts encoding important immune signaling components. For example, alternative splicing targets genes encoding PRRs, kinases, transcription factors, and NLRs^38–45^. In addition, the decapping and deadenylation protein complex as well as NMD factors have been shown to regulate stress-responsive transcripts^46–51^. Accordingly, these changes at the level of RBPs and transcripts contribute to plant immune responses against viruses (which depend on host translation) and other pathogens^37,50,51^.

ROS play an important role for biological processes such as plant development and responses to abiotic and biotic stresses, but are also extremely reactive and toxic at high levels, making its regulation critical to homeostasis^52^. Fine-tuning of ROS production and accumulation happens at different levels in space and time^52^, including post-translational modification of NAPDH oxidases. For instance, the most highly expressed NAPDH oxidase, RBOHD, is actively regulated to fine tune ROS production to permit growth, signaling and development while avoiding toxicity at high level^52–57^. Recently, post-translational modifications through phosphorylation and ubiquitination of RBOHD were shown to regulate its accumulation during immunity^57^. Our work here suggests that CBE1 and other translational regulators represent another layer of regulation of RBOHD protein accumulation; however, the mechanistic details underlying it remains unknown. This study nevertheless further emphasizes the importance of regulating ROS production through modulation of RBOHD. Investigating if CBE1 binds *RBOHD* transcripts directly, or binds other transcripts whose products regulate RBOHD levels, will be important to further understand the role of CBE1. In order to determine if this is part of a regulated attenuation mechanism, it will also be necessary to determine if RBOHD is under immune-induced translational control. Interestingly, recent results demonstrated that during immune signaling, *RBOHD* transcripts increased in the set of ribosome-loaded mRNAs^58^. However, the role of CBE1 in that process is still unknown, and expressing CBE1 in plants and bacteria has proven challenging^22^. Accordingly, we failed to generate stable Arabidopsis transgenic lines expressing epitope-tagged CBE1 despite multiple attempts (Table S2); highlighting the importance of generating novel tools to answer these questions in future studies.

Based on previous work showing the association between CBE1 and eIF4E1^22^, as well as the co-localization and mutant analysis presented here, we suggest that CBE1 might work together with decapping factor DCP1 and translation initiation factor eIF4E1 to regulate RBOHD protein level and consequently elicitor-induced ROS production and immunity. We found that mutants lacking initiation factor *eif4e* showed similar sensitivity to elf18 as *cbe1*, whereas mutants in other initiation factors (*eif4iso4e* and *eifiso4g1 eifiso4g2*) were similar to WT. These results are in accordance with the specificities of the different eIF isoforms, which bind the 5’ mRNA cap with a range of affinities^59,60^. We also observed enhanced elf18-induced ROS and RBOHD accumulation in *pat1-1*, which is surprising as eIF4E1 and PAT1 are predicted to function antagonistically. Indeed, eIF4E initiates recruitment of the initiation complex and subsequent recruitment of ribosomes, while PAT1 contributes to decapping, which initiates 5’-3’ decay by exoribonucleases^32^. In addition, CBE1 seems to localize predominantly to P-bodies, which are generally associated with mRNA decay^61^. Interestingly, the number of P-bodies increases when Arabidopsis is treated with flg22^32,50^, suggesting a link between P-body-mediated mRNA stability and immunity. Given that CBE1 is a plant-specific and non-essential protein, it has been proposed to regulate targeted transcripts in a context-dependent manner^22^, which could conceivably provide a fine-tuning mechanism to regulate gene expression. Further work is needed to understand how CBE1 functions in translation initiation and/or mRNA decay.

## Author contributions

J.G., M.S, J.M., and C.Z. conceived and designed the project. J.G., M.S, J.M. generated materials, performed experiments, and analyzed data. J.G. and C.Z. wrote the manuscript with input from all authors.

## Acknowledgments

This work was funded by The Gatsby Charitable Foundation (to C.Z.), The Biotechnology and Biological Research Council (BB/P012574/1), the University of Zürich (to C.Z.), and the Swiss National Science Foundation grant no. 31003A_182625 (to C.Z.). M.S. was supported by the Deutsche Forschungsgemeinschaft (Fellowship STE 2448/1 to M.S.), and J.M. by the European Molecular Biology Organization (Fellowships ALTF 459-2011). The authors thank Jonathan Jones (The Sainsbury Laboratory) for his input on the project, as well as Marta Bjornson, Julien Gronnier and all members of the Zipfel lab for fruitful discussion and feedback on the manuscript. The authors thank Julia Bailey-Serres for her input on the manuscript. The authors thank the John Innes Centre horticultural staff and Tamaryn Ellick (Institute of Plant and Microbial Biology) for assistance with plant growth. We also thank Karen Browning (The University of Texas at Austin) for providing the *cbe1-1, cum1-1, eif4g, eifiso4e, eifiso4g1 eifiso4g2* mutants; Morten Petersen (University of Copenhagen) for the *pat1-1, summ2-8, pat1-1 summ2-8* mutants; Martin Hülskamp (University of Cologne) for the *p35S-mCherry-PAB2, p35S-YFP-UBP1B, p35S-YFP-EIF4E1, p35S-YFP-RBP47C* constructs; and Martin Crespi (Institute of Plant Sciences Paris-Saclay) for the *p35S-mRFP1-UPF1, p35S-mRFP1-DCP1* constructs.

## STAR Methods text

### Plant materials and growth conditions

*Arabidopsis thaliana* plants were grown on soil as one to four plants per pot (7 × 7 cm) in controlled environment rooms maintained at 20 °C with a 10-h photoperiod and 60% humidity, or as seedlings on sterile Murashige and Skoog (MS) media supplemented with vitamins and 1%(w/v) sucrose (Duchefa) with a 16-h photoperiod. Assays using soil-grown plants were performed at 4 to 6 weeks post-germination (wpg), before the reproductive transition. Assays using plate-grown seedlings were performed at 2 wpg.

*A. thaliana* ecotype Columbia-0 (Col-0) was used as a wild-type control for all plant assays and was the background for all mutants used in this study, except otherwise stated. The *bak1-5 mob7* mutants were purified by one backcross to *bak1-5*. The single *mob7* mutant was obtained by crossing *bak1-5 mob7* to Col-0. Knock-down alleles *cbe1-2* (*AT4G01290*; SALK_038452), *cbe1-3* (*AT4G01290*; GK_150_H09), and wild-type alleles denoted *CBE1-2, CBE1-3* derived from segregation of SALK_038452 and GK_150_H09, respectively, were obtained through the Nottingham Arabidopsis Stock Centre (NASC). Ecotype Landsberg *erecta* (L*er-0*) and *rbohD* (SLAT line)^62^, *bak1-5* (EMS mutant)^23^ mutants were available in-house. Genotypes *cbe1-1* (WiscDsLoxHs188_10F)^22^, *eif4e1* (*cum1-1*)^63^, *eif4g* (SALK_80031)^22^, *eifiso4e* (SLAT line)^64^, double mutant *eifiso4g1 eifiso4g2* (from those two SALK lines: SALK_009905, SALK_076633)^65^ were obtained from Karen Browning. Genotypes *pat1-1* (SALK_040660), *summ2-8* (SAIL_1152A06), *pat1-1 summ2-8*^32^ were obtained from Morten Petersen.

*Nicotiana benthamiana* plants were grown on soil as one plant per pot (8 × 8 cm) at 25 °C during the day with 16 h light and at 22 °C during the night (8 h). Relative humidity was maintained at 60%.

### Map-based cloning and whole-genome sequencing

The *bak1-5 mob7* mutant (in Col-0) was crossed to L*er*-0. Fifty-six F_2_ segregants were genotyped for *bak1-5* using a dCAPS marker (Table S1). Homozygous *bak1-5* segregants were phenotyped for elf18-induced ROS production similar to *mob7*. Linkage analysis was performed using an array of genome-wide markers designed in-house or by the Arabidopsis Mapping Platform (Table S1)^66^. For whole-genome sequencing, 440 F_2_ plants from the cross *bak1-5 mob7* with *bak1-5* were scored for chitin-induced ROS production. One hundred thirty-three plants showed moderately increased, and 93 plants highly increased, ROS production. Out of these 93 plants, 70 were tested in the F_3_ generation, and only 15 showed a confirmed phenotype to restore Atpep1-induced seedling growth inhibition in 3 experiments. Thirty seedlings from each of the positive F_3_ parents were bulked and ground to a fine powder in liquid nitrogen and gDNA extracted. Ground tissues were equilibrated in buffer containing 50 mM Tris-HCl (pH 8.0), 200 mM NaCl, 2 mM EDTA for 30 min at 37 °C with occasional mixing, and a further 20 min at 37 °C with 0.2 mg/mL RNase. Roughly 10 ng of genomic DNA was then extracted using a standard chloroform/phenol method and resuspended in TE buffer (10 mM Tris HCl pH 7.5;1 mM EDTA pH 8). Prepared gDNA of pooled *bak1-5 mob7* F_3_ segregants, as well as *bak1-5* as a reference, was submitted to The Beijing Genomics Institute (Hong Kong) for Illumina-adapted library preparation and paired-end sequencing using the High-Seq 2000 platform. The average coverage from Illumina sequencing of *bak1-5 mob7* over the nuclear chromosomes was 15.79. Paired-end reads were aligned to the TAIR10 reference assembly using BWA v 0.6.1 with default settings^67^. BAM files were generated using SAMTools v 0.1.8^67^ and single-nucleotide polymorphisms (SNPs) were called using the mpileup command. High-quality SNPs were obtained using the filters (1) Reads with mapping quality less than 20 were ignored; (2) SNP position had a minimum coverage of 6 and a maximum of 250; (3) the reference base must be known; and (4) SNPs were present in *bak1-5 mob7* but not in the *bak1-5* control. The resulting pileup files contained a list of SNPs and their genomic positions. SNPs unique to *bak1-5 mob7* and not present in the *bak1-5* control were identified. SNPs passing filters were analyzed on CandiSNP^68^. Relevant SNPs were confirmed in the original *bak1-5 mob7* mutant and backcrossed lines by Sanger sequencing of PCR amplicons.

### Elicitors

The following elicitors were used in this study: chitin (Yaizu Suisankagaku Industry), flg22 peptide (CKANSFREDRNEDREV)^69^, elf18 peptide (ac-SKEKFERTKPHVNVGTIG)^70^, and Atpep1 peptide (ATKWKAKQRGKEKVSSGRPGQHN)^71^. All peptides were synthesized by SciLight-peptide (China) with purity above 95% and dissolved in sterile distilled water.

### Oxidative burst assay

Reactive oxygen species (ROS) production was measured as previously described^23^. For the assay, either adult plants (4- to 6-week-old plants) or seedlings (2-week-old) were used. For adult plants, leaf discs (4-mm diameter) were collected using a biopsy punch and floated overnight on distilled, deionized water in a white 96-well plate to recover from wounding. For ROS assays on whole seedlings, seedlings were grown on MS agar plates for 5 d before being transferred to MS liquid medium in transparent 96-well plates. After 8 d, seedlings were transferred to a white 96-well plate and allowed to recover overnight in sterile water. The water was then removed and replaced with elicitor solution containing 17 μg/mL luminol (Sigma-Aldrich), 100 μg/mL horseradish peroxidase (Sigma-Aldrich) and the indicated elicitor concentration. For seedlings, the hyperactive luminol derivative 0.5 μM L-012 (Fujifilm Wako Chemicals) was used instead of luminol. Luminescence was recorded over a 40- to 60-minute period using a charge-coupled device camera (Photek Ltd., East Sussex UK).

### Seedling growth inhibition assay

Seedling growth inhibition was performed as previously described^23^. Sterilized and stratified seeds were sown on MS media and grown in controlled environment rooms with 16/8 h day/night cycle and constant temperature of 22 °C. Five-day-old seedlings were transferred into liquid MS with or without the indicated amount of elicitor. Ten to twelve days later, individual seedlings were gently dry-blotted and weighed using a precision scale (Sartorius).

### MAP kinase phosphorylation assay

Phosphorylation of MAPKs was measured as previously described ^72^. Leaf discs (4-mm diameter) from adult plants (4- to 6-week-old plants) were cut in the evening and left overnight on the bench, floating in 6-well plates on distilled, deionized water. In the morning, the elicitor peptide was added to the desired concentration, and tissue was blotted dry and flash-frozen in liquid nitrogen for protein extraction at the indicated time points. MAPK phosphorylation was detected by western blot using an antibody specific to the active phosphorylated form of the proteins (phospho-p44/42 MAPK). Fifteen leaf discs were used per condition.

### Bacterial spray inoculation

Spray inoculations were performed as previously described^73^. *Pseudomonas syringae* pv. *tomato* (*Pto*) DC3000 wild-type and *COR*^*-*^ (defective in production of the phytotoxin coronatine) strains^74^ were grown in overnight culture in King’s B medium supplemented with 50 μg/mL rifampicin, 50 μg/mL kanamycin and 100 μg/mL spectinomycin and incubated at 28 °C. Cells were harvested by centrifugation and pellets resuspended in 10 mM MgCl_2_ to an OD_600_ of 0.2, corresponding to 1×10^8^ CFU/mL. Immediately before spraying, Silwet L-77 (Sigma Aldrich) was added to a final concentration of 0.04%(v/v). Four- to five-week-old plants were uniformly sprayed with the suspension and covered with a clear plastic lid for 3 d. Three leaf discs (4-mm diameter) were taken using a biopsy puncher from three respective leaves of one plant and ground in collection microtubes (Qiagen), containing one glass bead (3-mm diameter) and 200 μL water, using a 2010 Geno/Grinder (SPEX) at 1,500 rpm for 1.5 min. Ten microliters of serial dilutions from the extracts were plated on LB agar medium containing antibiotics and 25 μg/mL nystatin (Melford). Colonies were counted after incubation at 28°C for 1.5 to 2 d.

### Molecular cloning

Gateway-compatible fragments were amplified using Phusion Taq polymerase (New England Biolabs) from either Col-0 genomic DNA (*gCBE1*) containing 2.5 kb of the promoter sequence upstream of the translational start codon or from Col-0 complementary DNA (*cCBE1*) or from complementary *mob7* DNA (*cCBE1*^*mob7*^) and with or without the endogenous stop codon. attB flanked PCR products were cloned into pDONR201 using the BP Clonase II (Invitrogen) and recombination was performed using the LR Clonase II (Invitrogen) into the corresponding destination vector (pK7WGF2.0, pK7FWG2.0, pGWB604, pUBC-GFP-Dest, pB7WGR2.0)^75–77^. All clones were verified by Sanger sequencing.

### Transient expression in *N. benthamiana*

*N. benthamiana* plants were used for transient transformation at 4- to 5-week post-germination. *Agrobacterium tumefaciens* GV3101 overnight cultures grown at 28 °C in low-salt LB were harvested by centrifugation at 2,500 x *g* and resuspended in buffer containing 10 mM MgCl_2_ and 10 mM MES for 3 h at room temperature. *A. tumefaciens*-mediated transient transformation of *N. benthamiana* was performed by infiltrating leaves with OD_600_ = 0.2 of each construct together with the viral suppressor P19 in a 1:1 (or 1:1:1) ratio. Samples were collected 2-3 d after infiltration.

### Stable transformation of Arabidopsis

Transgenic Arabidopsis plants were generated using floral dip method^78^. Briefly, flowering Arabidopsis plants were dipped into suspension culture of *Agrobacterium tumefaciens* GV3101 carrying the indicated plasmid. Plants carrying a T-DNA insertion event were selected either on MS medium containing the appropriate selection or as soil-grown seedlings by spray application of Basta (Bayer Crop Science).

### Confocal microscopy

*N. benthamiana* leaf discs (4-mm diameter) transiently over-expressing the indicated proteins were sampled at 2-3 dpi with water as the imaging medium. Live-cell imaging employed a laser-scanning Leica SP5 Confocal Microscope (Leica Microsystems, Wetzlar, Germany) and 63x (glycerol immersion) objective. GFP was excited at 488 nm and emission detected between 496–536 nm (shown in green). YFP was excited at 514 nm and detected between 524–551 nm (shown in yellow). RFP derivatives (mRFP, mCherry, tag-RFP) were excited at 561 nm and detected between 571-635 nm (shown in magenta). Co-localization was performed using sequential channel analysis by calculating Pearson’s coefficient^25,79^ using the coloc 2 plugin of ImageJ. Image analysis was performed with Fiji.

### Western blotting

Antibodies used in western blots were as follows: anti-GFP 1:5000; Santa Cruz); α-RFP-HRP (1:5000; Abcam); anti-mouse-HRP (1:15000; Sigma Aldrich); anti-rabbit-HRP (1:10000; Sigma Aldrich); anti-RBOHD (1:1000; Agrisera) and anti-phospho-p42/p44-erk (1:1000; Cell Signalling Tech).

### Statistical analysis

Statistical analysis was performed using R (4.1.2) and Rstudio (2021.09.1) or GraphPad Prism (9.3). Based on Gaussian distribution parametric or nonparametric tests were chosen and when n ≥30, normal distribution was assumed. Prior to multiple comparisons, ANOVA or Kruskal-Wallis test were performed to look for differences across groups. For multiple comparisons, Dunnett’s and Dunn’s tests were favored to compare multiple groups to one control group. Tests were realized on the overall set of replicates and replicates were included only when positive and negative controls showed the expected results.

## Supplemental item titles

**Figure S1.**
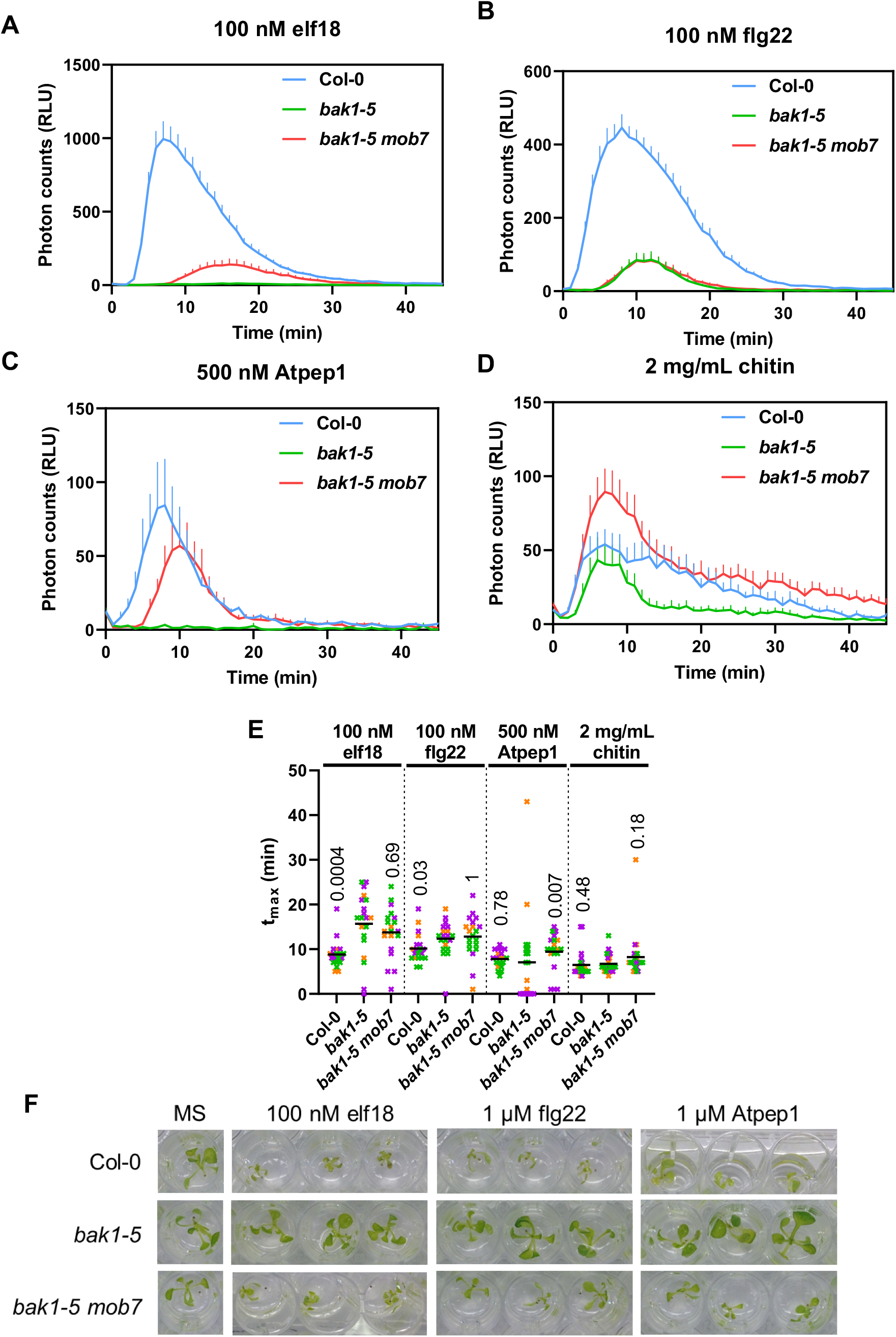
*mob7* restores immune signalling in *bak1-5*. Related to Figure 1. (A-D) ROS burst kinetic measured as relative light units (RLU), in leaf discs following treatment with 100 nM elf18 (A) or 100 nM flg22 (B) or 500 nM Atpep1 (C) or 2 mg/mL chitin (D). Values are means + standard errors (n=8). (E) Tmax describes the time it takes for the ROS to peak upon treatment with corresponding elicitors over 60 min recording.. Horizontal lines represent the means from 3 independent experiments (n=4-8). The symbol colors indicate the different experiments. Numbers above symbols are p-values from Dunn’s multiple comparison test between corresponding genotype and *bak1-5*. (F) Images of 14-day-old seedlings grown in MS media or MS media containing 100 nM elf18, 1 μM flg22 or 1 μM Atpep1.

**Figure S2.**
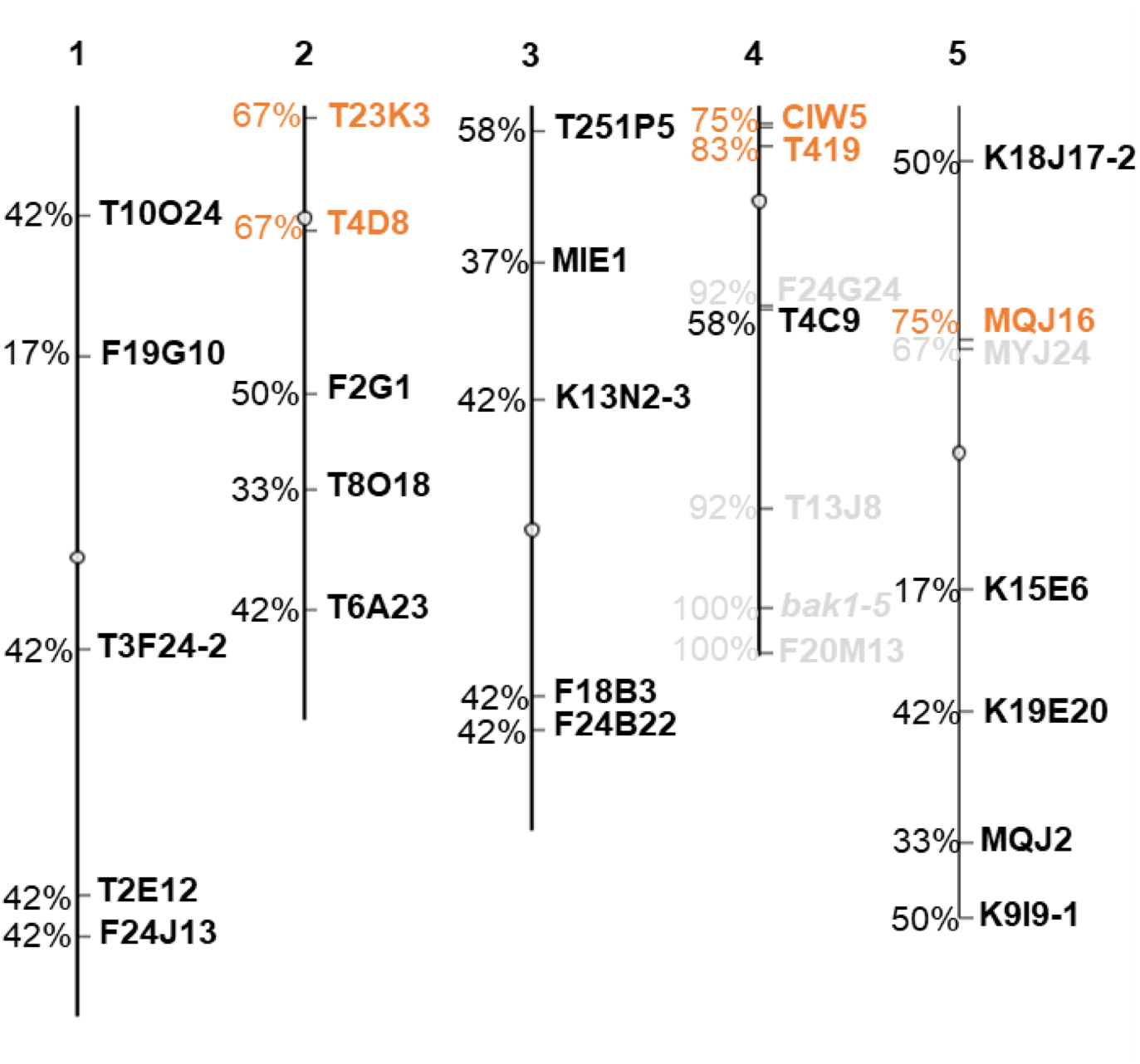
Map-based cloning of *bak1-5 mob7*. Physical linkage map constructed using the F_2_ population from *bak1-5 mob7* x Ler-0. The percentage represent the percentage of Col-0 alleles contributed by *bak1-5 mob7*. Plants were screened based on ROS production upon treatment with 100 nM elf18. Markers in grey are markers for which an increase of Col-0 alleles was also observed in plants with low elicitor-induced ROS production, thereby removed from further analysis. Markers in orange show linkage statistically higher than 50%. Circles represent centromeres. Significance was determined by χ^2^ test.

**Figure S3.**
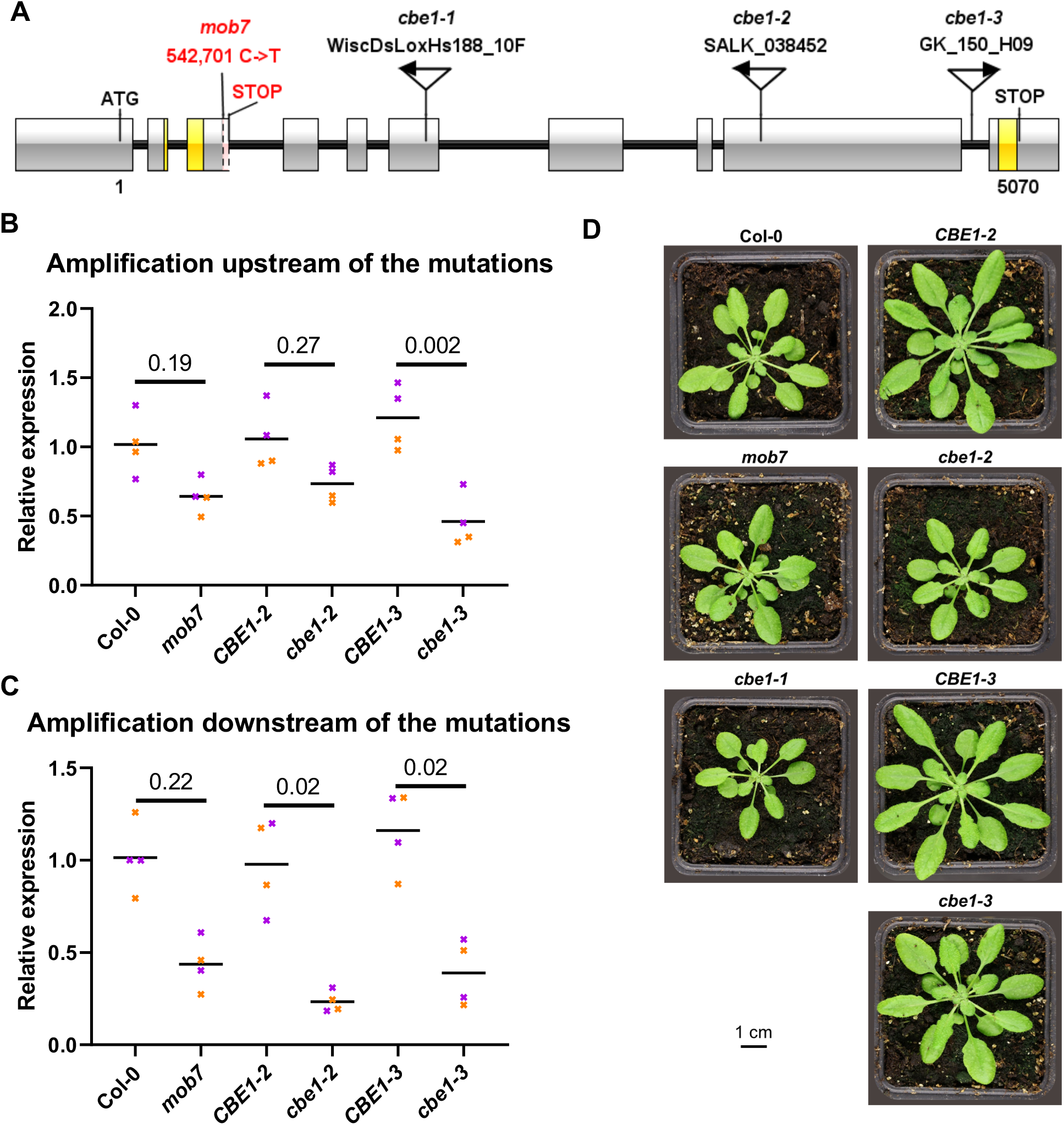
Characterization of *cbe1* alleles. (A) Gene structure of *AT4G01290*. Exons are shown as grey boxes. T-DNA insertions are indicated with the respective name above and arrows indicate the orientation of the T-DNAs. EMS-induced *mob7* mutation with respective position and early stop codon are indicated in red. Fragments amplified by quantitative reverse-transcription polymerase chain reaction (RT-qPCR) are indicated in yellow. Bp, base pairs. (B, C) RT-qPCR of *AT4G01290* upstream of the T-DNA insertions/*mob7* mutation (B) or downstream of the insertions/mutation (C). (B,C) Expression values relative to the *U-BOX* housekeeping gene are shown. CBE1-2 and CBE1-3 are *CBE1* wildtype segregants from the *cbe1-2* and *cbe1-3* lines, respectively. Horizontal lines represent the means from 2 independent experiments (n=2) (B,C) The symbol colours indicate the different experiments. Numbers above horizontal lines are p-values from Dunn’s multiple comparison test between genotypes under the lines. (D) Rosette morphology of 5-week-old plants of the corresponding genotype.

**Figure S4.**
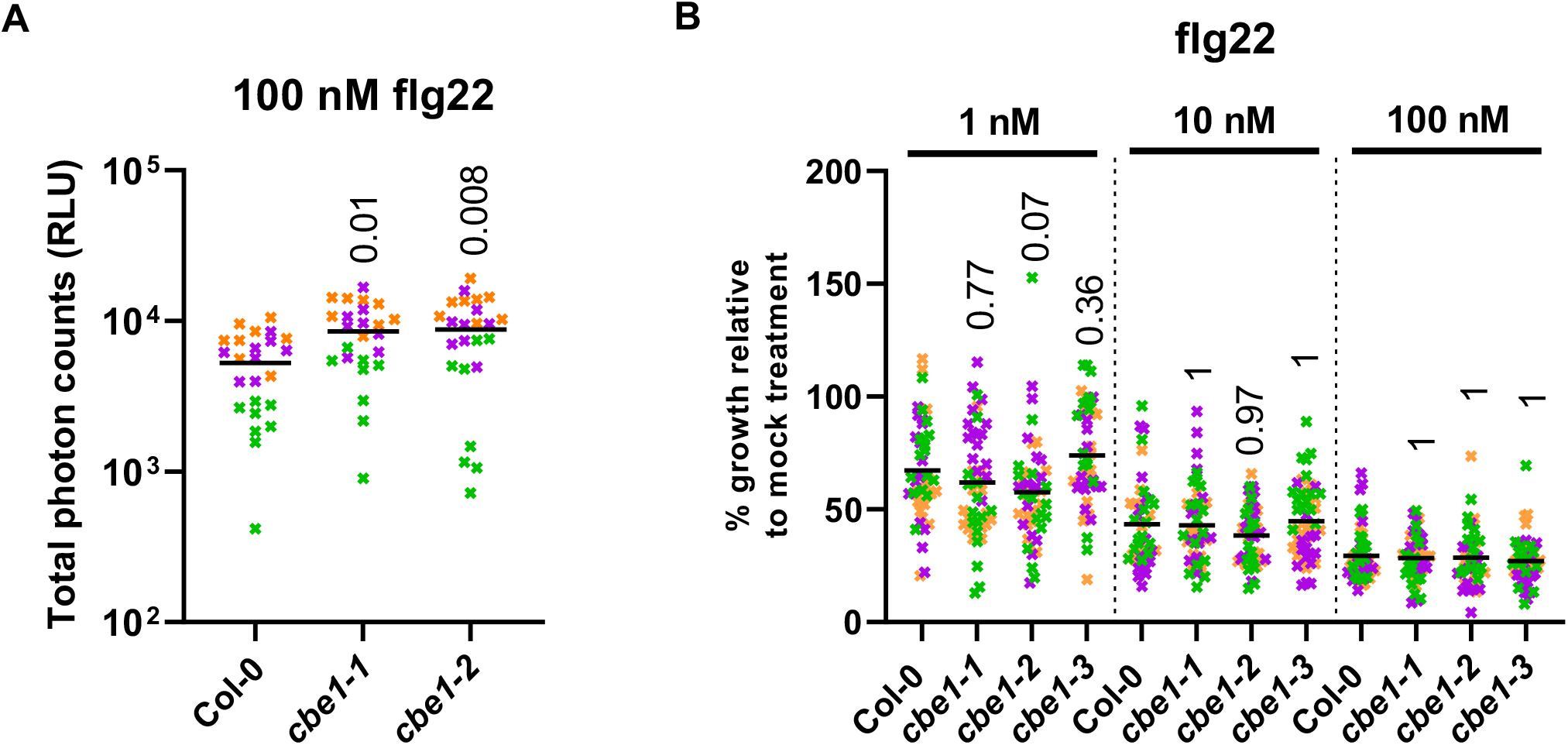
CBE1 negatively regulates elicitor-induced ROS production. (A) Total ROS accumulation measured as RLU over 60 min recording after treatment with 100 nM flg22 on leaf discs from 5-week-old plants: Horizontal lines represent the means from 3 independent experiments (n=8). (B) Bacterial growth (CFU/cm^2^) in leaves spray inoculated with 10^7^ CFU/mL (OD_600_ =0.2) *P. syringae* pv. *tomato* DC3000 *COR*^*-*^ and sampled at 3 dpi. Horizontal lines represent the means from 4 independent experiments (n=7-8). (C) Growth inhibition represented as percentage of fresh weight in response to 1, 10 or 100 nM flg22 relative to mock treated seedlings. Horizontal lines represent the means from 3 independent experiments (n=16). Numbers above symbols are p-values from (A,B) Dunnett’s or (C) Dunn’s multiple comparison test between corresponding genotype and *bak1-5*.

**Figure S5.**
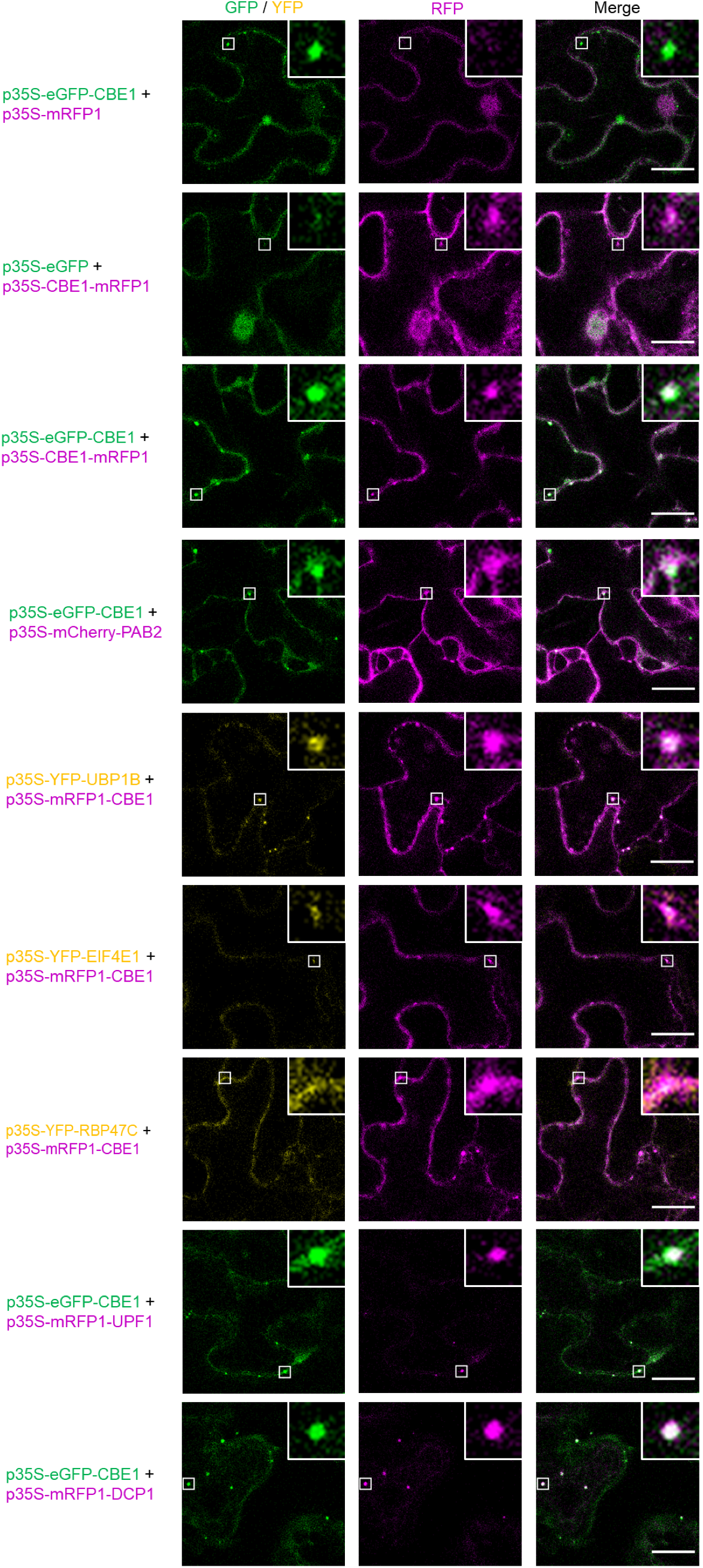
CBE1 localizes predominantly to processing bodies among ribonucleoprotein complexes. Confocal images of green, yellow and red fluorescent proteins. The proteins were transiently co-expressed in *N. benthamiana*. Confocal microscopy on leaf discs was conducted 3 days post-infiltration. Merged pictures show overlay of GFP/YFP and RFP. The scale bar corresponds to 20 μm. An ROI of 25 μm^2^ is shown by white square and zoomed in on the top right of the images. P-bodies markers: UPF1, DCP1; polysomes/stress granules markers: PAB2, EIF4E; stress granules markers: UBP1B, RBP47C.

**Table S1.**
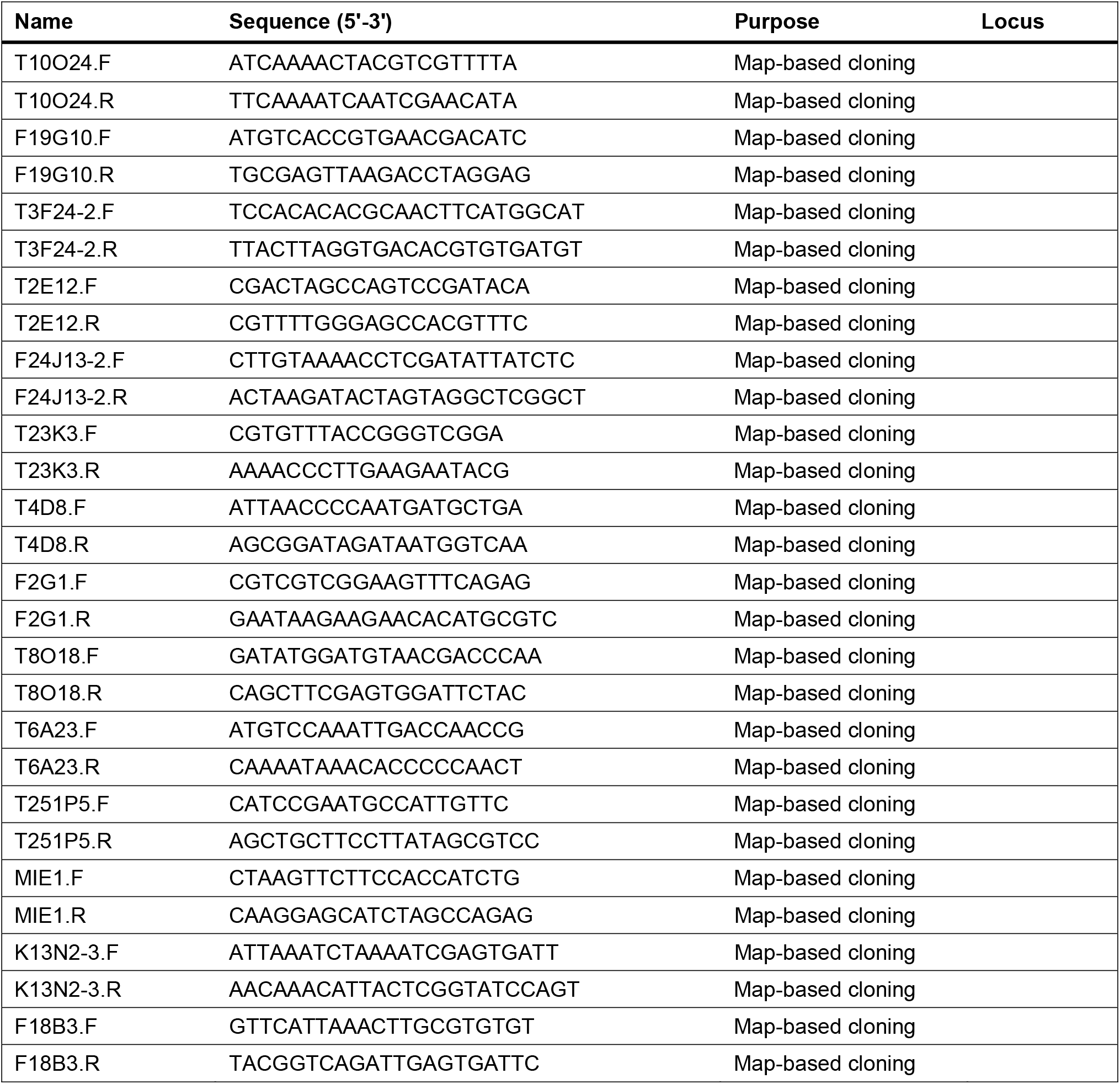
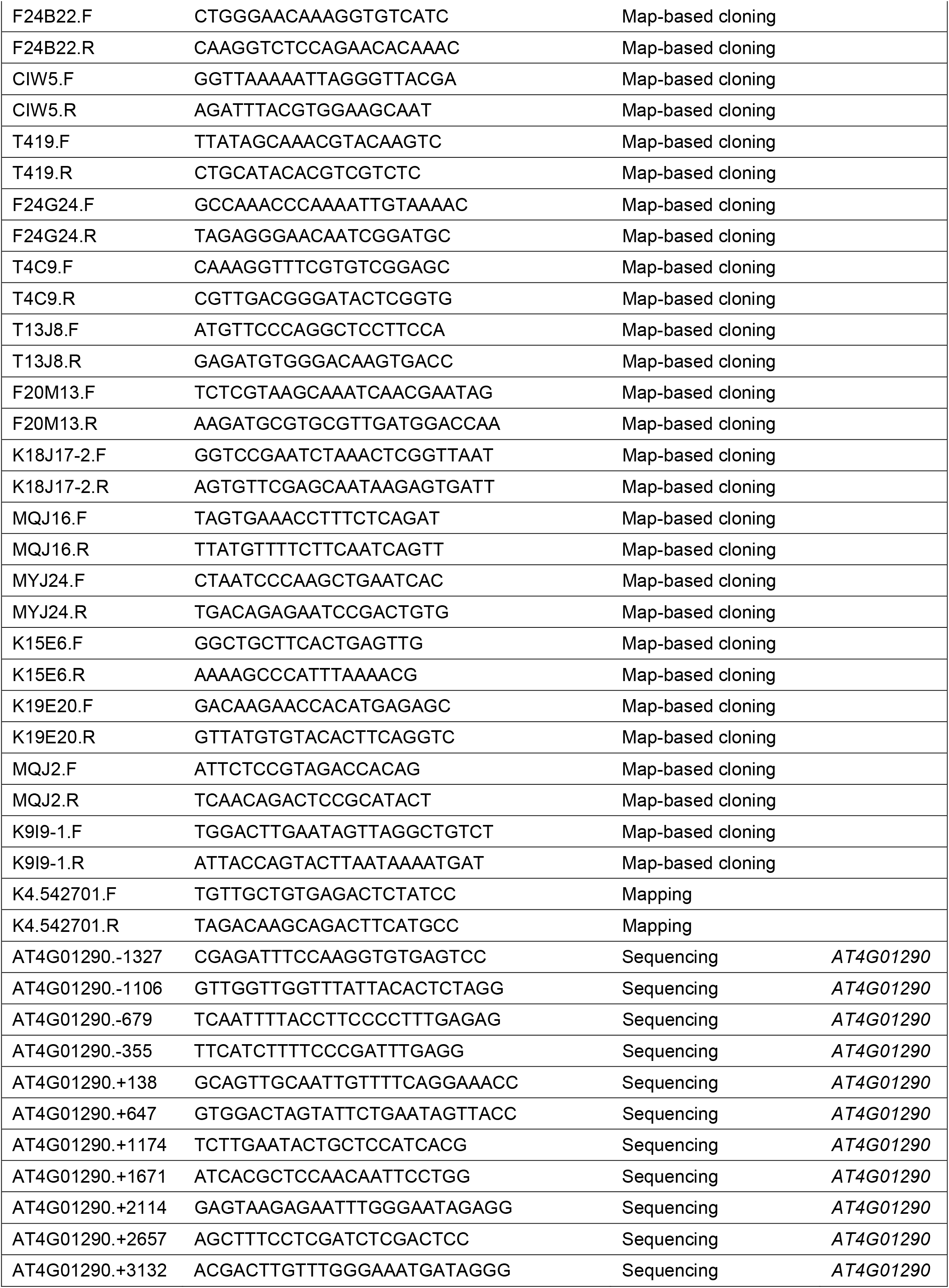
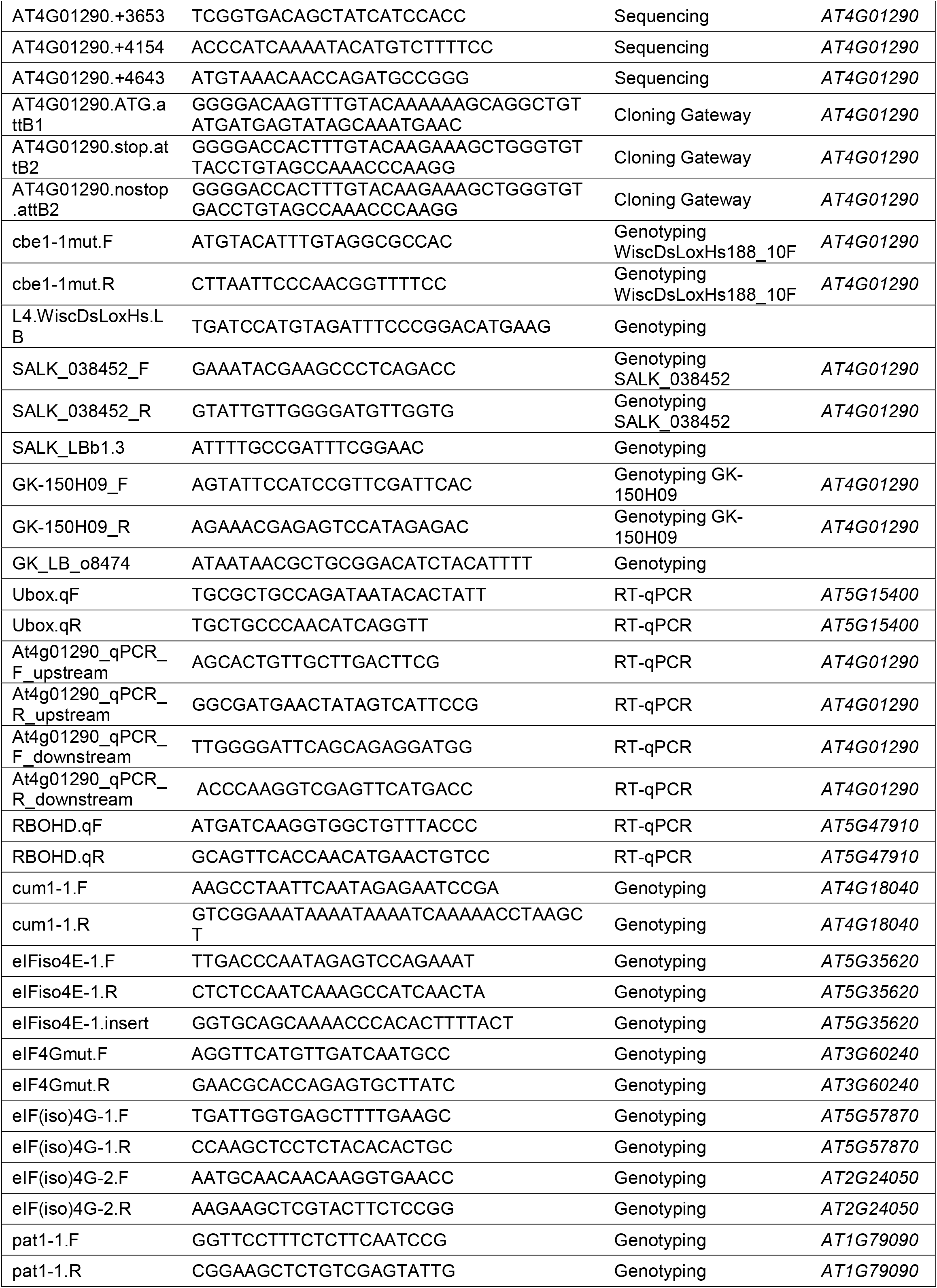
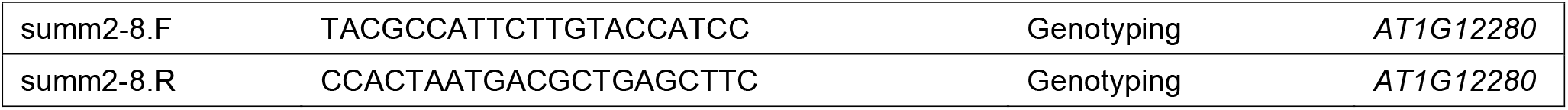
Primers used in this study.

**Table S2.** *CBE1* transgenics investigated.

| Genetic background | Construct | Backbone | Outcome |
| --- | --- | --- | --- |
| Col-0 | *p35S-eGFP-cCBE1* | pK7WGF2.0 | No expression of the transgene |
| Col-0 | *p35S-cCBE1-eGFP* | pK7FWG2.0 | No expression of the transgene |
| Col-0 | *pCBE1-gCBE1-eGFP* | pGWB604 | No expression of the transgene |
| Col-0 | *pUBI10-cCBE1-eGFP* | pUBC-GFP-Dest | No expression of the transgene |
| *bak1-5 mob7* | *p35S-eGFP-cCBE1* | pK7WGF2.0 | No expression of the transgene |
| *bak1-5 mob7* | *p35S-cCBE1-eGFP* | pK7FWG2.0 | No expression of the transgene |
| *bak1-5 mob7* | *pCBE1-gCBE1-eGFP* | pGWB604 | No expression of the transgene |
| *bak1-5 mob7* | *pUBI10-cCBE1-eGFP* | pUBC-GFP-Dest | No expression of the transgene |
| *cbe1-1* | *p35S-eGFP-cCBE1* | pK7WGF2.0 | No expression of the transgene |
| *cbe1-1* | *p35S-cCBE1-eGFP* | pK7FWG2.0 | No expression of the transgene |
| *cbe1-1* | *pCBE1-gCBE1-eGFP* | pGWB604 | No expression of the transgene |
| *cbe1-1* | *pUBI10-cCBE1-eGFP* | pUBC-GFP-Dest | No expression of the transgene |

## References

1. Gust, A.A., Pruitt, R., and Nürnberger, T. (2017). Sensing Danger: Key to Activating Plant Immunity. Trends Plant Sci. 22, 779–791.

2. Boller, T., and Felix, G. (2009). A renaissance of elicitors: Perception of microbe-associated molecular patterns and danger signals by pattern-recognition receptors. Annu. Rev. Plant Biol. 60, 379–407.

3. Boutrot, F., and Zipfel, C. (2017). Function, Discovery, and Exploitation of Plant Pattern Recognition Receptors for Broad-Spectrum Disease Resistance. Annu. Rev. Phytopathol. 55, 257–286.

4. Gómez-Gómez, L., and Boller, T. (2000). FLS2: an LRR receptor-like kinase involved in the perception of the bacterial elicitor flagellin in Arabidopsis. Mol. Cell 5, 1003–11.

5. Zipfel, C., Kunze, G., Chinchilla, D., Caniard, A., Jones, J.D.G., Boller, T., and Felix, G. (2006). Perception of the bacterial PAMP EF-Tu by the receptor EFR restricts Agrobacterium-mediated transformation. Cell 125, 749–60.

6. Yamaguchi, Y., Pearce, G., and Ryan, C.A. (2006). The cell surface leucine-rich repeat receptor for AtPep1, an endogenous peptide elicitor in Arabidopsis, is functional in transgenic tobacco cells. Proc. Natl. Acad. Sci. U. S. A. 103, 10104–9.

7. Chinchilla, D., Zipfel, C., Robatzek, S., Kemmerling, B., Nürnberger, T., Jones, J.D.G., Felix, G., and Boller, T. (2007). A flagellin-induced complex of the receptor FLS2 and BAK1 initiates plant defence. Nature 448, 497–500.

8. Heese, A., Hann, D.R., Gimenez-Ibanez, S., Jones, A.M.E., He, K., Li, J., Schroeder, J.I., Peck, S.C., and Rathjen, J.P. (2007). The receptor-like kinase SERK3/BAK1 is a central regulator of innate immunity in plants. Proc. Natl. Acad. Sci. U. S. A. 104, 12217–22.

9. Roux, M., Schwessinger, B., Albrecht, C., Chinchilla, D., Jones, A., Holton, N., Malinovsky, F.G., Tör, M., de Vries, S., and Zipfel, C. (2011). The Arabidopsis leucine-rich repeat receptor-like kinases BAK1/SERK3 and BKK1/SERK4 are required for innate immunity to hemibiotrophic and biotrophic pathogens. Plant Cell 23, 2440–55.

10. Yu, X., Feng, B., He, P., and Shan, L. (2017). From Chaos to Harmony: Responses and Signaling upon Microbial Pattern Recognition. Annu. Rev. Phytopathol. 55, 109–137.

11. DeFalco, T.A., and Zipfel, C. (2021). Molecular mechanisms of early plant pattern-triggered immune signaling. Mol. Cell 81, 3449–3467.

12. Monaghan, J., Matschi, S., Shorinola, O., Rovenich, H., Matei, A., Segonzac, C., Malinovsky, F.G., Rathjen, J.P., MacLean, D., Romeis, T., et al. (2014). The calcium-dependent protein kinase CPK28 buffers plant immunity and regulates BIK1 turnover. Cell Host Microbe 16, 605–15.

13. Holmes, D.R., Bredow, M., Thor, K., Pascetta, S.A., Sementchoukova, I., Siegel, K.R., Zipfel, C., and Monaghan, J. (2021). A novel allele of the Arabidopsis thaliana MACPF protein CAD1 results in deregulated immune signaling. Genetics 217.

14. Stegmann, M., Monaghan, J., Smakowska-Luzan, E., Rovenich, H., Lehner, A., Holton, N., Belkhadir, Y., and Zipfel, C. (2017). The receptor kinase FER is a RALF-regulated scaffold controlling plant immune signaling. Science 355, 287–289.

15. Monaghan, J., Matschi, S., Romeis, T., and Zipfel, C. (2015). The calcium-dependent protein kinase CPK28 negatively regulates the BIK1-mediated PAMP-induced calcium burst. Plant Signal. Behav. 10, e1018497.

16. Wang, J., Grubb, L.E., Wang, J., Liang, X., Li, L., Gao, C., Ma, M., Feng, F., Li, M., Li, L., et al. (2018). A Regulatory Module Controlling Homeostasis of a Plant Immune Kinase. Mol. Cell 69, 493-504.e6.

17. Morita-Yamamuro, C., Tsutsui, T., Sato, M., Yoshioka, H., Tamaoki, M., Ogawa, D., Matsuura, H., Yoshihara, T., Ikeda, A., Uyeda, I., et al. (2005). The Arabidopsis gene CAD1 controls programmed cell death in the plant immune system and encodes a protein containing a MACPF domain. Plant Cell Physiol. 46, 902–12.

18. Chen, T., Nomura, K., Wang, X., Sohrabi, R., Xu, J., Yao, L., Paasch, B.C., Ma, L., Kremer, J., Cheng, Y., et al. (2020). A plant genetic network for preventing dysbiosis in the phyllosphere. Nature 580, 653–657.

19. Xiao, Y., Stegmann, M., Han, Z., DeFalco, T.A., Parys, K., Xu, L., Belkhadir, Y., Zipfel, C., and Chai, J. (2019). Mechanisms of RALF peptide perception by a heterotypic receptor complex. Nature 572, 270–274.

20. Gronnier, J., Franck, C.M., Stegmann, M., DeFalco, T.A., Abarca, A., Von Arx, M., Dünser, K., Lin, W., Yang, Z., Kleine-Vehn, J., et al. (2022). Regulation of immune receptor kinase plasma membrane nanoscale organization by a plant peptide hormone and its receptors. Elife 11, 2020.07.20.212233.

21. Bush, M.S., Hutchins, A.P., Jones, A.M.E., Naldrett, M.J., Jarmolowski, A., Lloyd, C.W., and Doonan, J.H. (2009). Selective recruitment of proteins to 5′ cap complexes during the growth cycle in Arabidopsis. Plant J. 59, 400–412.

22. Patrick, R.M., Lee, J.C.H., Teetsel, J.R.J., Yang, S., Choy, G.S., and Browning, K.S. (2018). Discovery and characterization of conserved binding of eIF4E 1 (CBE1), a eukaryotic translation initiation factor 4E-binding plant protein. J. Biol. Chem. 293, 17240–17247.

23. Schwessinger, B., Roux, M., Kadota, Y., Ntoukakis, V., Sklenar, J., Jones, A., and Zipfel, C. (2011). Phosphorylation-dependent differential regulation of plant growth, cell death, and innate immunity by the regulatory receptor-like kinase BAK1. PLoS Genet. 7, e1002046.

24. Brogna, S., and Wen, J. (2009). Nonsense-mediated mRNA decay (NMD) mechanisms. Nat. Struct. Mol. Biol. 16, 107–13.

25. Adler, J., and Parmryd, I. (2010). Quantifying colocalization by correlation: the Pearson correlation coefficient is superior to the Mander’s overlap coefficient. Cytometry. A 77, 733–42.

26. Chantarachot, T., and Bailey-Serres, J. (2018). Polysomes, Stress Granules, and Processing Bodies: A Dynamic Triumvirate Controlling Cytoplasmic mRNA Fate and Function. Plant Physiol. 176, 254–269.

27. Xu, J., Yang, J.-Y., Niu, Q.-W., and Chua, N.-H. (2006). Arabidopsis DCP2, DCP1, and VARICOSE Form a Decapping Complex Required for Postembryonic Development. Plant Cell 18, 3386–3398.

28. Kerényi, F., Wawer, I., Sikorski, P.J., Kufel, J., and Silhavy, D. (2013). Phosphorylation of the N- and C-terminal UPF1 domains plays a critical role in plant nonsense-mediated mRNA decay. Plant J. 76, 836–48.

29. Weber, C., Nover, L., and Fauth, M. (2008). Plant stress granules and mRNA processing bodies are distinct from heat stress granules. Plant J. 56, 517–530.

30. Sorenson, R., and Bailey-Serres, J. (2014). Selective mRNA sequestration by OLIGOURIDYLATE-BINDING PROTEIN 1 contributes to translational control during hypoxia in Arabidopsis. Proc. Natl. Acad. Sci. 111, 2373–2378.

31. Browning, K.S., and Bailey-Serres, J. (2015). Mechanism of Cytoplasmic mRNA Translation. Arab. B. 13, e0176.

32. Roux, M.E., Rasmussen, M.W., Palma, K., Lolle, S., Regué, À.M., Bethke, G., Glazebrook, J., Zhang, W., Sieburth, L., Larsen, M.R., et al. (2015). The mRNA decay factor PAT1 functions in a pathway including MAP kinase 4 and immune receptor SUMM2. EMBO J. 34, 593–608.

33. Kong, L., Rodrigues, B., Kim, J.H., He, P., and Shan, L. (2021). More than an on-and-off switch: Post-translational modifications of plant pattern recognition receptor complexes. Curr. Opin. Plant Biol. 63, 102051.

34. Meteignier, L.-V., El Oirdi, M., Cohen, M., Barff, T., Matteau, D., Lucier, J.-F., Rodrigue, S., Jacques, P.-E., Yoshioka, K., and Moffett, P. (2017). Translatome analysis of an NB-LRR immune response identifies important contributors to plant immunity in Arabidopsis. J. Exp. Bot. 68, 2333–2344.

35. Xu, G., Greene, G.H., Yoo, H., Liu, L., Marqués, J., Motley, J., and Dong, X. (2017). Global translational reprogramming is a fundamental layer of immune regulation in plants. Nature 545, 487–490.

36. Yoo, H., Greene, G.H., Yuan, M., Xu, G., Burton, D., Liu, L., Marqués, J., and Dong, X. (2020). Translational Regulation of Metabolic Dynamics during Effector-Triggered Immunity. Mol. Plant 13, 88–98.

37. Bach-pages, M., Chen, H., Sanguankiattichai, N., Soldan, R., Kaiser, M., Mohammed, S., Hoorn, R.A.L. Van Der, Castello, A., Preston, G.M., Kaschani, F., et al. (2020). Proteome-wide profiling of RNA-binding protein responses to flg22 reveals novel components of plant immunity. bioRxiv.

38. Dinesh-Kumar, S.P., and Baker, B.J. (2000). Alternatively spliced N resistance gene transcripts: their possible role in tobacco mosaic virus resistance. Proc. Natl. Acad. Sci. U. S. A. 97, 1908–13.

39. Zhang, X.-C., and Gassmann, W. (2007). Alternative splicing and mRNA levels of the disease resistance gene RPS4 are induced during defense responses. Plant Physiol. 145, 1577–87.

40. Gassmann, W. (2008). Alternative splicing in plant defense. Curr. Top. Microbiol. Immunol. 326, 219–33.

41. Yang, S., Tang, F., and Zhu, H. (2014). Alternative splicing in plant immunity. Int. J. Mol. Sci. 15, 10424–45.

42. Zhang, Z., Liu, Y., Ding, P., Li, Y., Kong, Q., and Zhang, Y. (2014). Splicing of receptor-like kinase-encoding SNC4 and CERK1 is regulated by two conserved splicing factors that are required for plant immunity. Mol. Plant 7, 1766–75.

43. Liu, J., Chen, X., Liang, X., Zhou, X., Yang, F., Liu, J., He, S.Y., and Guo, Z. (2016). Alternative Splicing of Rice WRKY62 and WRKY76 Transcription Factor Genes in Pathogen Defense. Plant Physiol. 171, 1427–42.

44. Bazin, J., Mariappan, K., Jiang, Y., Blein, T., Voelz, R., Crespi, M., and Hirt, H. (2020). Role of MPK4 in pathogen-associated molecular pattern-triggered alternative splicing in Arabidopsis. PLoS Pathog. 16, e1008401.

45. Dressano, K., Weckwerth, P.R., Poretsky, E., Takahashi, Y., Villarreal, C., Shen, Z., Schroeder, J.I., Briggs, S.P., and Huffaker, A. (2020). Dynamic regulation of Pep-induced immunity through post-translational control of defence transcript splicing. Nat. Plants.

46. Liang, W., Li, C., Liu, F., Jiang, H., Li, S., Sun, J., Wu, X., and Li, C. (2009). The Arabidopsis homologs of CCR4-associated factor 1 show mRNA deadenylation activity and play a role in plant defence responses. Cell Res. 19, 307–16.

47. Walley, J.W., Kelley, D.R., Savchenko, T., and Dehesh, K. (2010). Investigating the function of CAF1 deadenylases during plant stress responses. Plant Signal. Behav. 5, 802–5.

48. Gloggnitzer, J., Akimcheva, S., Srinivasan, A., Kusenda, B., Riehs, N., Stampfl, H., Bautor, J., Dekrout, B., Jonak, C., Jiménez-Gómez, J.M., et al. (2014). Nonsense-mediated mRNA decay modulates immune receptor levels to regulate plant antibacterial defense. Cell Host Microbe 16, 376–90.

49. Tabassum, N., Eschen-Lippold, L., Athmer, B., Baruah, M., Brode, M., Maldonado-Bonilla, L.D., Hoehenwarter, W., Hause, G., Scheel, D., and Lee, J. (2020). Phosphorylation-dependent control of an RNA granule-localized protein that fine-tunes defence gene expression at a post-transcriptional level. Plant J. 101, 1023–1039.

50. Yu, X., Li, B., Jang, G.-J., Jiang, S., Jiang, D., Jang, J.-C., Wu, S.-H., Shan, L., and He, P. (2019). Orchestration of Processing Body Dynamics and mRNA Decay in Arabidopsis Immunity. Cell Rep. 28, 2194-2205.e6.

51. Chantarachot, T., Sorenson, R.S., Hummel, M., Ke, H., Kettenburg, A.T., Chen, D., Aiyetiwa, K., Dehesh, K., Eulgem, T., Sieburth, L.E., et al. (2020). DHH1/DDX6-like RNA helicases maintain ephemeral half-lives of stress-response mRNAs. Nat. Plants 6, 675–685.

52. Castro, B., Citterico, M., Kimura, S., Stevens, D.M., Wrzaczek, M., and Coaker, G. (2021). Stress-induced reactive oxygen species compartmentalization, perception and signalling. Nat. plants 7, 403–412.

53. Chen, D., Cao, Y., Li, H., Kim, D., Ahsan, N., Thelen, J., and Stacey, G. (2017). Extracellular ATP elicits DORN1-mediated RBOHD phosphorylation to regulate stomatal aperture. Nat. Commun. 8, 2265.

54. Kadota, Y., Sklenar, J., Derbyshire, P., Stransfeld, L., Asai, S., Ntoukakis, V., Jones, J.D., Shirasu, K., Menke, F., Jones, A., et al. (2014). Direct regulation of the NADPH oxidase RBOHD by the PRR-associated kinase BIK1 during plant immunity. Mol. Cell 54, 43–55.

55. Li, L., Li, M., Yu, L., Zhou, Z., Liang, X., Liu, Z., Cai, G., Gao, L., Zhang, X., Wang, Y., et al. (2014). The FLS2-associated kinase BIK1 directly phosphorylates the NADPH oxidase RbohD to control plant immunity. Cell Host Microbe 15, 329–38.

56. Zhang, M., Chiang, Y.-H., Toruño, T.Y., Lee, D., Ma, M., Liang, X., Lal, N.K., Lemos, M., Lu, Y.-J., Ma, S., et al. (2018). The MAP4 Kinase SIK1 Ensures Robust Extracellular ROS Burst and Antibacterial Immunity in Plants. Cell Host Microbe 24, 379-391.e5.

57. Lee, D., Lal, N.K., Lin, Z.D., Ma, S., Liu, J., Castro, B., Toruño, T., Dinesh-Kumar, S.P., and Coaker, G. (2020). Regulation of reactive oxygen species during plant immunity through phosphorylation and ubiquitination of RBOHD. Nat. Commun. 11, 1838.

58. Ngou, B.P.M., Ahn, H.K., Ding, P., and Jones, J.D.G. (2021). Mutual potentiation of plant immunity by cell-surface and intracellular receptors. Nature 592, 110–115.

59. Khan, M.A., and Goss, D.J. (2018). Kinetic analyses of phosphorylated and non-phosphorylated eIFiso4E binding to mRNA cap analogues. Int. J. Biol. Macromol. 106, 387–395.

60. Kropiwnicka, A., Kuchta, K., Lukaszewicz, M., Kowalska, J., Jemielity, J., Ginalski, K., Darzynkiewicz, E., and Zuberek, J. (2015). Five eIF4E isoforms from Arabidopsis thaliana are characterized by distinct features of cap analogs binding. Biochem. Biophys. Res. Commun. 456, 47–52.

61. Luo, Y., Na, Z., and Slavoff, S.A. (2018). P-Bodies: Composition, Properties, and Functions. Biochemistry 57, 2424–2431.

62. Torres, M.A., Dangl, J.L., and Jones, J.D.G. (2002). Arabidopsis gp91phox homologues AtrbohD and AtrbohF are required for accumulation of reactive oxygen intermediates in the plant defense response. Proc. Natl. Acad. Sci. U. S. A. 99, 517–22.

63. Yoshii, M., Yoshioka, N., Ishikawa, M., and Naito, S. (1998). Isolation of an Arabidopsis thaliana mutant in which accumulation of cucumber mosaic virus coat protein is delayed. Plant J. 13, 211–219.

64. Duprat, A., Caranta, C., Revers, F., Menand, B., Browning, K.S., and Robaglia, C. (2002). The Arabidopsis eukaryotic initiation factor (iso)4E is dispensable for plant growth but required for susceptibility to potyviruses. Plant J. 32, 927–34.

65. Nicaise, V., Gallois, J.-L., Chafiai, F., Allen, L.M., Schurdi-Levraud, V., Browning, K.S., Candresse, T., Caranta, C., Le Gall, O., and German-Retana, S. (2007). Coordinated and selective recruitment of eIF4E and eIF4G factors for potyvirus infection in Arabidopsis thaliana. FEBS Lett. 581, 1041–6.

66. Hou, X., Li, L., Peng, Z., Wei, B., Tang, S., Ding, M., Liu, J., Zhang, F., Zhao, Y., Gu, H., et al. (2010). A platform of high-density INDEL/CAPS markers for map-based cloning in Arabidopsis. Plant J. 63, 880–8.

67. Li, H., and Durbin, R. (2009). Fast and accurate short read alignment with Burrows-Wheeler transform. Bioinformatics 25, 1754–60.

68. Etherington, G.J., Monaghan, J., Zipfel, C., and MacLean, D. (2014). Mapping mutations in plant genomes with the user-friendly web application CandiSNP. Plant Methods 10, 41.

69. Felix, G., Duran, J.D., Volko, S., and Boller, T. (1999). Plants have a sensitive perception system for the most conserved domain of bacterial flagellin. Plant J. 18, 265–76.

70. Kunze, G., Zipfel, C., Robatzek, S., Niehaus, K., Boller, T., and Felix, G. (2004). The N terminus of bacterial elongation factor Tu elicits innate immunity in Arabidopsis plants. Plant Cell 16, 3496–507.

71. Huffaker, A., Pearce, G., and Ryan, C. a (2006). An endogenous peptide signal in Arabidopsis activates components of the innate immune response. Proc. Natl. Acad. Sci. U. S. A. 103, 10098–103.

72. Flury, P., Klauser, D., Schulze, B., Boller, T., and Bartels, S. (2013). The anticipation of danger: microbe-associated molecular pattern perception enhances AtPep-triggered oxidative burst. Plant Physiol. 161, 2023–35.

73. Katagiri, F., Thilmony, R., and He, S.Y. (2002). The Arabidopsis thaliana-pseudomonas syringae interaction. Arab. B. 1, e0039.

74. Melotto, M., Underwood, W., Koczan, J., Nomura, K., and He, S.Y. (2006). Plant stomata function in innate immunity against bacterial invasion. Cell 126, 969–80.

75. Grefen, C., Donald, N., Hashimoto, K., Kudla, J., Schumacher, K., and Blatt, M.R. (2010). A ubiquitin-10 promoter-based vector set for fluorescent protein tagging facilitates temporal stability and native protein distribution in transient and stable expression studies. Plant J. 64, 355–65.

76. Karimi, M., Inzé, D., and Depicker, A. (2002). GATEWAY vectors for Agrobacterium-mediated plant transformation. Trends Plant Sci. 7, 193–5.

77. Nakamura, S., Mano, S., Tanaka, Y., Ohnishi, M., Nakamori, C., Araki, M., Niwa, T., Nishimura, M., Kaminaka, H., Nakagawa, T., et al. (2010). Gateway binary vectors with the bialaphos resistance gene, bar, as a selection marker for plant transformation. Biosci. Biotechnol. Biochem. 74, 1315–9.

78. Clough, S.J., and Bent, A.F. (1998). Floral dip: a simplified method for Agrobacterium-mediated transformation of Arabidopsis thaliana. Plant J. 16, 735–43.

79. Dunn, K.W., Kamocka, M.M., and McDonald, J.H. (2011). A practical guide to evaluating colocalization in biological microscopy. Am. J. Physiol. Cell Physiol. 300, C723–42.

